# Peroxisomal Import Stress Drives Cellular Senescence through SCAF1-dependent Suppression of Mitoribosome Biogenesis

**DOI:** 10.64898/2026.09.19.752713

**Authors:** Jacinta Correia, Jinoh Kim, Pham Thuy Tien Vo, Hua Bai

## Abstract

Communication between peroxisomes and mitochondria is essential for cellular metabolic homeostasis, yet how peroxisomal import stress impacts mitochondria function during aging and cellular senescence remains poorly defined. Using a genome-wide CRISPR screening in HEK293 cells under peroxisome import stress, we identified SCAF1 (SR-related CTD-associated factor 1) a known canonical nuclear pre-mRNA splicing factor, as an essential regulator of mitochondrial homeostasis. Under peroxisome stress SCAF1 undergoes proteolytic processing and translocates to the mitochondria, where its N-terminal region acts as an autonomous repressor module that blocks mitoribosomal subunit joining. Consequently, SCAF1 depletion accelerates subunit joining and elevates oxidative phosphorylation protein levels, whereas its overexpression in IMR90 fibroblast cells triggers robust cellular senescence characterized by increased senescence associated β gal staining. Together, our findings uncover a stress-responsive peroxisome-to-mitochondria signaling axis mediated by SCAF1 translocation. This pathway directly modulates mitoribosome assembly to maintain translational homeostasis, providing a precise molecular mechanism for how upstream peroxisomal decline drives downstream mitochondrial dysfunction and cellular senescence.

## INTRODUCTION

Aging is accompanied by a progressive decline in cellular homeostasis (*1*, *2*) driven in part by dysfunction of energy metabolizing organelles and impaired coordination between them(*3*, *4*). Mitochondria play a central role in ATP production, redox signaling, apoptosis, and metabolic control (*4*, *5*), and their dysfunction is a hallmark of aging and cellular senescence (*1*, *6–8*). Increasing evidence indicates that age-associated mitochondrial decline does not occur in isolation but instead reflects failures in inter-organelle communication pathways that coordinate metabolism, stress responses, and protein homeostasis across compartments (*9*, *10*). Among these networks, the cross-talk between peroxisomes and mitochondria are metabolically coupled organelles that jointly execute fatty acid β-oxidation, reactive oxygen species (ROS) management, lipid remodeling, and cellular redox regulation (*11–16*). Peroxisomes initiate β-oxidation of very-long chain fatty acids (VLCFAs), synthesize plasmalogen, and perform ether phospholipid biosynthesis and cellular redox regulation functions that depend entirely on the import of nuclear-encoded matrix proteins through a set of conserved import receptors called peroxins (PEXs) (*12*, *17*, *18*). The PEX5 receptor delivers PTS1-bearing cargo into the peroxisomal matrix and is recycled to the cytosol through a monoubiquitination extraction cycle that requires a conserved N-terminal cysteine (Cys11 in human PEX5) (*19–22*). Disruption of peroxisomal import, as occurs through genetic mutations in PEX genes in peroxisome biogenesis disorders (PBDs) such as Zellweger spectrum disorder, leads to severe impairment of fatty acid oxidation and plasmalogen biosynthesis and, ultimately, profound neurological dysfunction (*14*, *17*, *18*). Recent work from our group has further shown that peroxisomal import declines with age in *Drosophila* oenocytes, human B cells and tissues, linking organelle dysfunction to systemic inflammation and age-related pathologies (*23–25*). Despite their fundamental importance, peroxisomes remain among the least mechanistically characterized of the major metabolic organelles, and the molecular responses that cells deploy to cope with peroxisomal dysfunction remain incompletely defined.

Peroxisomal dysfunction has direct and well-documented consequences for mitochondrial function. Hepatocyte-specific *Pex5* knockout mice exhibit altered mitochondria structure, reduced oxidative phosphorylation (OXPHOS) activity, and elevated mitochondrial ROS production (*26*, *27*). Similar secondary mitochondrial dysfunction, including impaired respiratory chain complex activity and altered mitochondrial morphology, has been documented in fibroblasts from Zellweger syndrome patients and in additional *Pex* mutant model systems (*13*, *27*). Peroxisomes and mitochondria share not only metabolic substrates but also fission machinery components, including DRP1 and FIS1 (*28*), and maintain physical proximity through dedicated membrane contact sites involving the peroxisomal tethering factor ACBD5 (*29–31*) and the mitochondrial outer membrane scaffold TOMM20 (*28*, *29*). Despite these well-established structural and metabolic connections, the molecular mechanisms by which peroxisomal dysfunction is actively sensed and transduced into adaptive changes in mitochondrial gene expression remain entirely unknown.

A central node of mitochondrial gene expression is the biogenesis and assembly of the mitoribosome, a highly specialized ribonucleoprotein complex responsible for translating the thirteen mitochondrially encoded subunits of the OXPHOS complexes (*32–39*). Unlike cytosolic ribosomes, the human mitoribosome is membrane-anchored at the inner mitochondrial membrane and consists of a 39S large subunit (mt-LSU) and a 28S small subunit (mt-SSU) that join to form a 55S monosome (*32–34*, *40*, *41*). Mitoribosome assembly requires an elaborate cohort of nuclear-encoded factors, including DEAD-box and DEAH-box RNA helicases (DDX28, DHX30), RNA-binding proteins (LRPPRC, SLIRP, FASTKD4), methyltransferases (MRM2, MRM3), and a sequential GTPase-driven maturation pathway involving GTPBP5, GTPBP6, GTPBP7, and GTPBP10 (*40*, *42–53*). Late-stage maturation of the mt-LSU is gated by the MALSU1·L0R8F8·mt-ACP anti-association module, which physically prevents premature 28S–39S subunit joining at the intersubunit bridge (*37*, *48*, *54*, *55*). Many of these assembly steps occur within mitochondrial RNA granules nucleoid-proximal, membraneless compartments that concentrate nascent rRNA, mRNA, and assembly factors and that mediate the earliest steps of 16S rRNA maturation and ribosome biogenesis (*38*, *50*, *56*, *57*). Disruption of any step of this pathway impairs mitochondrial translation, compromises respiratory chain stoichiometry, and triggers mitochondrial dysfunction (*58–60*). Despite the complexity and importance of this pathway, no stress-responsive mechanism has been described by which an extrinsic signal from a dysfunctional organelle can arrest mitoribosome assembly to adaptively regulate translational output.

SR (serine/arginine-rich) proteins constitute a major family of nuclear RNA-binding factors defined by their RS-repeat domains and roles in pre-mRNA splicing, mRNA export, mRNA stability, and translational regulation (*61–66*). Several SR proteins shuttle between the nucleus and cytoplasm in a phosphorylation-dependent manner regulated by SRPK and CLK family kinases, allowing them to participate in cytoplasmic RNA-regulatory processes beyond their canonical nuclear splicing functions (*65*, *66*). SCAF1 (SR-Related CTD Associated Factor 1) is a member of this family, interacting with the C-terminal domain of RNA Polymerase II and contributing to pre-mRNA splicing and transcriptional co-regulation and polyadenylation site usage (*67–71*). SCAF1 is overexpressed in aggressive ovarian and breast cancers (*67*, *68*), has recently been identified as a tumour suppressor in pancreatic ductal adenocarcinoma in a recent in vivo CRISPR screen (*70*), and is genetically associated with cardiac conduction disorders (*72*), another identified individuals with loss-of function variants in SCAF1 typically show cognitive impairment like autism (*73*). Despite these disease associations, the biological function of SCAF1 remains largely unknown. Although the splicing factor PRP-19 has been linked to the mitochondrial unfolded protein response (*74*), that role is exerted through nuclear splicing rather than direct mitochondrial localization; no SR-related CTD -associated splicing factor has been shown to be imported into mitochondria or to act directly within organelle.

An emerging theme in cellular stress biology is the deployment of canonical nuclear factors to non-canonical organelle-protective roles under stress, a paradigm of protein moonlighting that is now well established for metabolic enzymes, RNA-binding proteins, and splicing factors. (*74*– *77*) The mitochondrial unfolded protein response (mtUPR), exemplifies this bidirectional organelle-nucleus communication and was originally characterized in *C. elegans* through the transcription factor ATFS-1 (*78*, *79*) and subsequently extended to mammalian cells through ATF5 (*80*, *81*), illustrates the broader principle that mitochondrial proteostasis is actively communicated to and from the nucleus. (*82*, *83*) Nuclear-encoded mitochondrial proteins are imported into the organelle, while mitochondrial status (e.g., proteotoxic load) feeds back to regulate nuclear gene expression, creating a feedback loop that co-ordinates biogenesis, quality control and repair. Distinct mitochondrial defects engage the cytosolic integrated stress response (ISR) through the OMA1–DELE1–HRI relay axis,(*84–88*) coupling mitochondrial perturbations to translational reprogramming. However, no analogous mechanism has been identified that conveys peroxisomal status to the mitochondrial gene expression machinery.

Crucially, dysregulated mitochondrial translation (*89*) and peroxisomal dysfunction (*10*) have been recently linked to aging and cellular senescence, which exhibit elevated ROS, chronic inflammation, and a senescence associated secretory phenotype (SASP) that drives tissue dysfunction (*8*). However, the direct molecular axis connecting these events has remained missing. In this study we demonstrate that peroxisomal stress or forced mitochondrial targeting of SCAF1 is sufficient to induce senescence markers, positioning SCAF1-mediated mitochondrial translational control as a key determinant of cell fate decisions under metabolic stress. Understanding SCAF1-mediated signaling will illuminate mechanisms underlying age-associated mitochondrial dysfunction and cellular senescence. In the long term, modulating SCAF1 activity or mitochondrial translation may provide new strategies to delay senescence, restore organelle homeostasis, or ameliorate age-related diseases.

## RESULTS

### Genome-wide CRISPR screening identifies SCAF1 as a novel regulator of peroxisomal stress response

Although peroxisome dysfunction is increasingly recognized as a driver of lipid metabolic dysregulation and broader organelle dysfunction, the cellular machinery that detects defective peroxisomes and coordinates cytoprotective responses has yet to be defined. To address this gap, we engineered a tetracycline-inducible system of peroxisomal stress in HEK293 cells, enabling systematic identification of the key molecular components underlying the cellular response. A Tet-On 3G expression construct encoding FLAG-tagged human PEX5 carrying a cysteine-to-alanine substitution at position 11 (PEX5^C11A^) was knocked into the AAVS1 safe-harbour locus (*25*). Cysteine 11 is the conserved monoubiquitination site required for PEX5 recycling, and its substitution produces a dominant-negative mutant that blocks peroxisomal matrix protein import and recapitulates the molecular basis of Zellweger spectrum disorder (*19–22*, *25*). Robust induction of PEX5^C11A^ was achieved by treatment with 1μg/ml doxycycline (Dox), that causes a sharp reduction of peroxisomal import within 24hrs.

To comprehensively map which genes are involved in regulating cell fitness under peroxisomal stress, we performed a pooled CRISPR-Cas9 knockout screening using Brunello lentiviral sgRNA library (*90*, *91*)that contains 76,441 sgRNAs for 19,114 human genes (about 4 sgRNAs per gene) in Tet-PEX5^C11A^ HEK293 cells (**Fig. 1A**). The library was transduced into Cas9-expressing *Tet-PEX5^C11A^* knock-in HEK293 cells. Seven days later, peroxisomal stress was induced with doxycycline. After three rounds of treatment, genomic DNA was isolated and sgRNA abundance compared between Dox-treated (Dox+) and untreated (Dox-) cells by NGS (**Fig. 3A**). This identified 1,528 genes significantly enriched or depleted (fold change >2, p<0.05) (**Fig. 1B**), with many hits in spliceosome, RNA degradation, ribosome, metabolic pathway, oxidative phosphorylation (OXPHOS) (**Fig. 1C**). This finding aligns with the well-established functional interplay between peroxisomes and mitochondria [10], as well as our recent discovery linking peroxisomes to cytosolic ribosome biogenesis [20].

**Fig 1.**
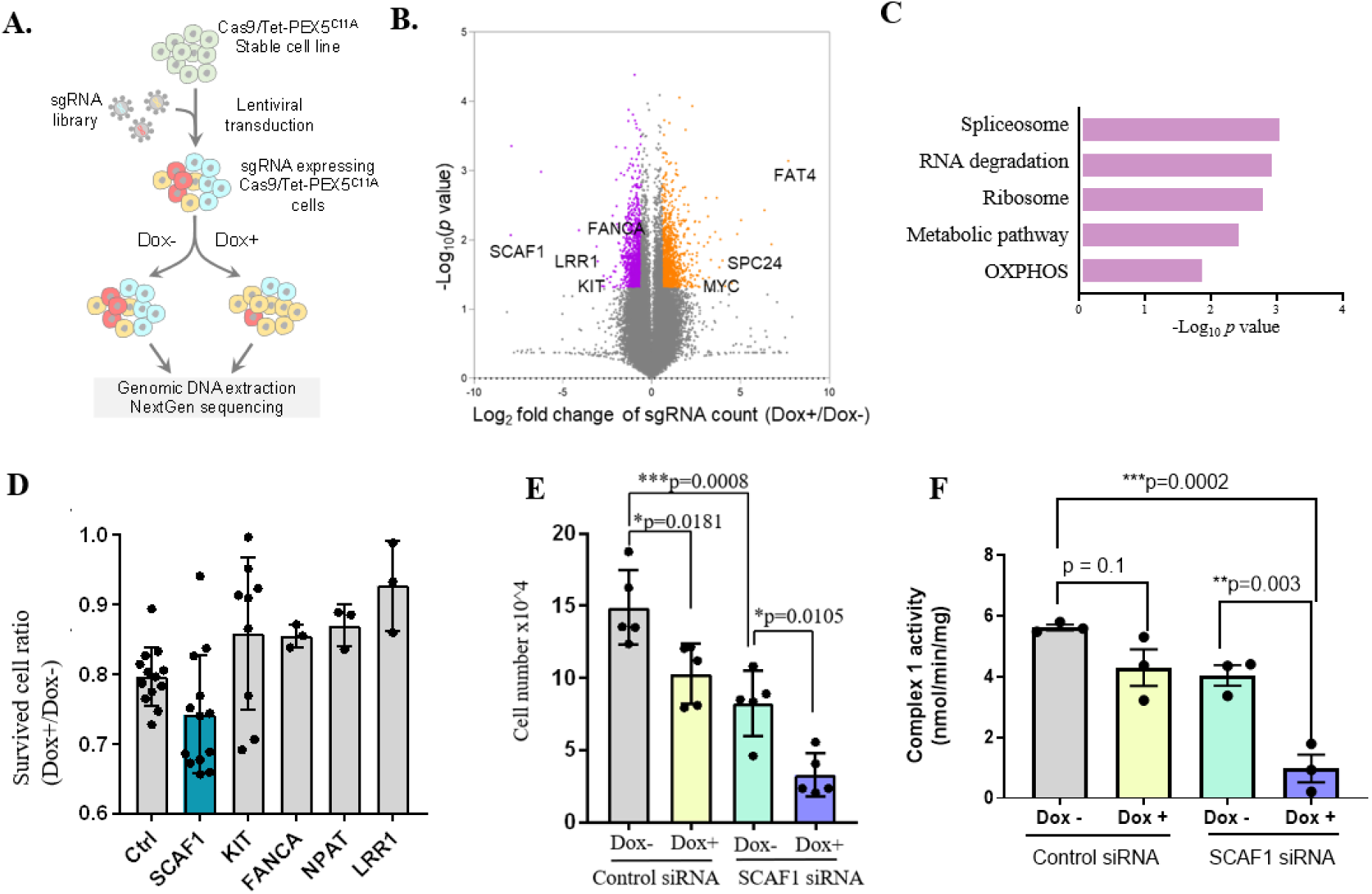
SCAF1 is identified as a key modifier to maintain cellular fitness upon peroxisomal import stress. A)Schematic diagram for genome-wide CRISPR knockout screening using Tet-On PEX5^C11A^ cells w/wo doxycycline treatment (1μg/ml). (B) Volcano plot showing differentially enriched sgRNAs and top target genes from CRISPR knockout screening. (C) Gene Ontology (GO) enrichment analysis of top candidate pathways. (D) Validation of candidate genes by cell survival assay. Tet-On PEX5^C11A^ cells were transfected with control siRNA or gene-specific siRNAs targeting candidate genes (*SCAF1, KIT, FANCA, NPAT, LRR1*) w/wo doxycycline treatment (1μg/ml). After 2 days, surviving cells were counted, and the ratio of doxycycline-treated (Dox+) to untreated (Dox-) cells were calculated. mean ± SD. (E) Cell number was counted after knockdown SCAF1 for 3 days w/wo doxycycline treatment (1 μg/ml). One-way ANOVA test, data presented as mean ± SD. (F) Mitochondrial Complex I activity ( nmol oxidized NADH/mom/mg protein) in Tet-PEX5^C11A^ cells transfected with scramble or SCAF1 siRNA under Dox-induced peroxisomal stress. N = 3. Two-way ANOVA. *, *p*<0.05. ns, not significant

**Fig. 2.**
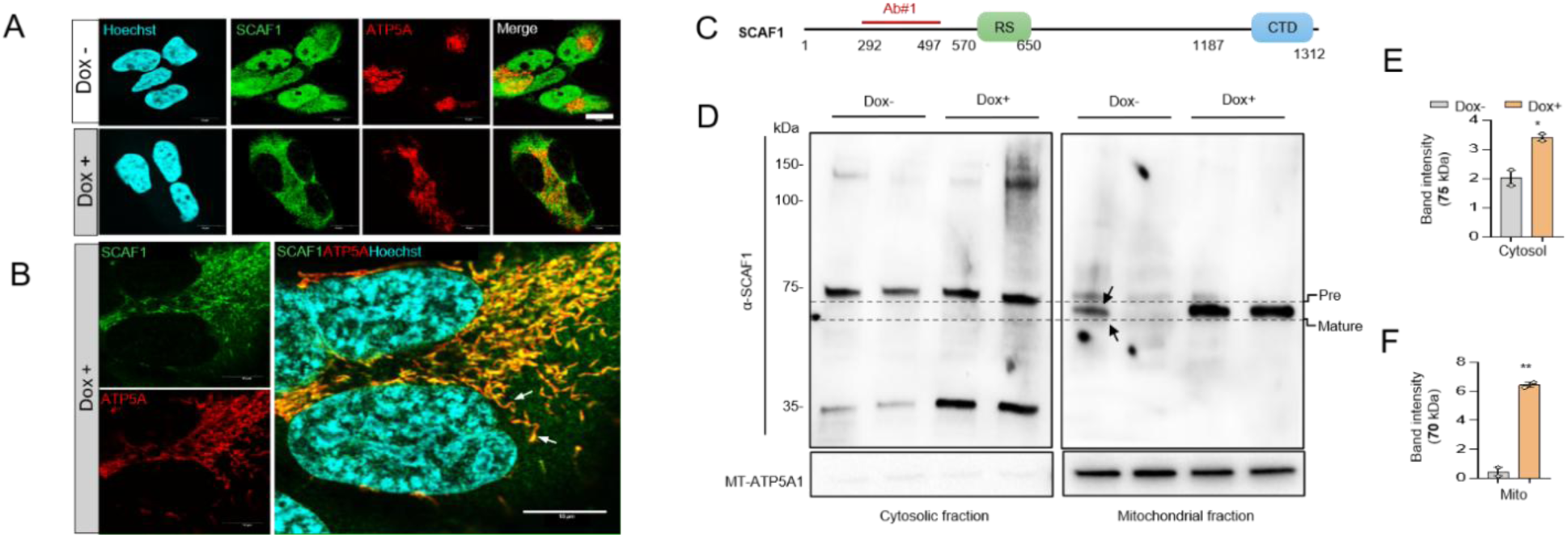
Peroxisomal stress induces mitochondrial translocation and processing of SCAF1. (A) Immunofluorescence images showing enhanced mitochondrial localization of SCAF1 under peroxisomal stress (Dox+), compared with predominant nuclear localization under basal conditions (Dox-). ATP5A was used as a mitochondrial marker. Scale bar: 10 μm. (B) Higher-magnification images highlighting stress-induced mitochondrial translocation of SCAF1 (white arrows). (C) Schematic representation of the SCAF1 protein, indicating the serine/arginine-rich (RS) domain and C-terminal domain (CTD); the antibody immunogen region is highlighted in red. (D) Immunoblotting of subcellular fractions revealing a distinct SCAF1 isoform (black arrows) that accumulates in mitochondria upon peroxisomal stress (Dox+). (E, F) Quantification of the cytosolic SCAF1 precursor (75 kDa) and the processed mitochondrial form (70 kDa). N = 2, t-test. *, *p*<0.05. **, *p*<0.01.

**Fig 3.**
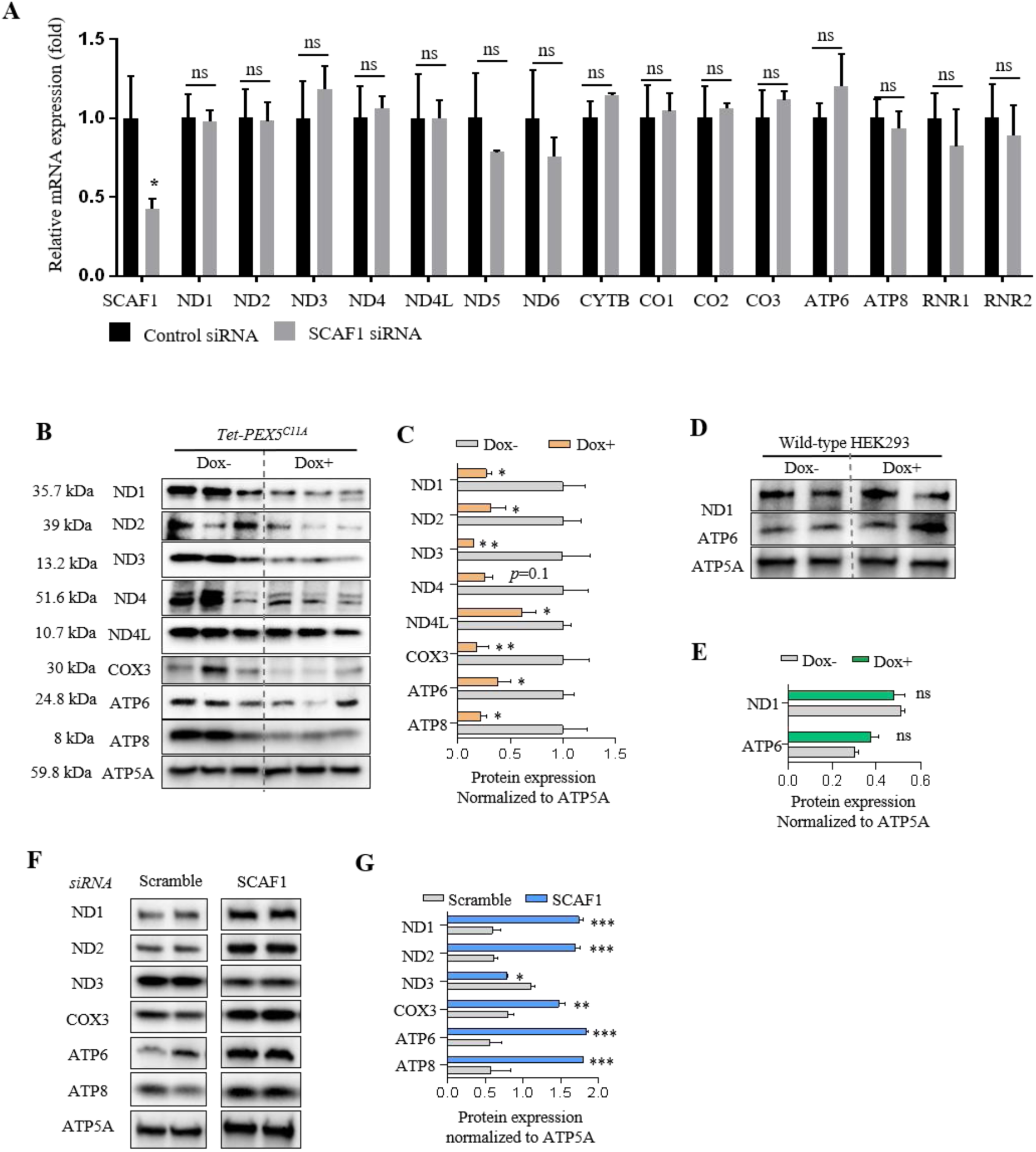
SCAF1 represses mitochondrial translation under peroxisomal stress. (A) qRT-PCR analysis of mtDNA-encoded transcript levels following SCAF1 KD. N = 3; Student’s t-test. (B) Immunoblotting analysis of mtDNA-encoded proteins in cells subjected to peroxisomal stress (Dox+, 1 μg/mL). ATP5A is a mitochondrial loading control. (C) Quantification of protein levels shown in panel B. N = 3. (D) Immunoblotting analysis of mtDNA-encoded proteins in wild-type cells treated with Dox (1 μg/mL) alone. (E) Quantification of protein levels shown in panel D. N = 2. (F) Immunoblotting analysis of mtDNA-encoded proteins following SCAF1 KD. (G) Quantification of protein levels shown in panel F. N = 2; Student’s t-test. *, p<0.05. **, p<0.01. ***, p<0.001.

To validate the screening results, we performed targeted siRNA knockdowns of top candidates (*SCAF1*, *KIT*, *FANCA*, *NPAT*, and *LRR1*) and measured cell survival ratios (Dox+/Dox−) after three days of induced stress. *SCAF1* knockdown resulted in the most severe reduction in cellular fitness among all candidates tested (**Fig. 1D**). Absolute cell counting confirmed that while SCAF1 depletion alone moderately impacted basal growth, the combination of *SCAF1* knockdown and Dox-induced peroxisomal stress synergistically impaired cell proliferation and survival (**Fig. 1E**), establishing. Consistent with a role as a stress-responsive regulator, we found that both *SCAF1* mRNA (**Fig. S1.G**) and SCAF1 protein (**Fig. S1. H**) levels were robustly upregulated in parallel with Dox-induced PEX5^C11A^ expression.

To define the broader transcriptomic landscape of this stress response, we performed RNA-sequencing and Gene Set Enrichment Analysis (GSEA). Three functionally coherent pathway responses emerged: a severe downregulation of OXPHOS/electron transport chain pathways, a coordinated suppression of mitochondrial ribosome biogenesis/translation, and a marked upregulation of the Integrated Stress Response (ISR) and Unfolded Protein Response (UPR) (**Fig. S1. I**). SCAF1 is a serine/arginine-rich (SR) pre-mRNA splicing factor [16] whose C-terminal domain interacts with the CTD of RNA polymerase II, implicating it in pre-mRNA splicing [23] and the serine/arginine-rich (RS) domain is thoughts to be required for protein-protein interactions [24]. Its broader biological function remains largely unknown, though it is highly expressed in ovarian and breast cancers [16, 25, 26] and was recently identified as a driver of pancreatic cancer [27]. Interestingly, *SCAF1* KD significantly induced mitochondrial dysfunction under peroxisomal stress, as indicated by impaired Complex I activity (**Fig. 1F**). Together, our findings reveal an unexpected role for splicing factor SCAF1 in peroxisome-mitochondria crosstalk.

### SR family splice factor SCAF1 translocates to mitochondria under peroxisome stress

Having established that SCAF1 is required for cellular fitness under peroxisomal import stress, we next sought to determine how a canonical nuclear splicing factor (*61*, *67–69*) involved with mitochondria. Therefore, we next asked where SCAF1 localizes when peroxisomal function is compromised. We examined the subcellular localization of SCAF1 to determine its functional sites. Using an in-house SCAF1 antibody (Booster Bione DZ41740; **Fig.2.C**), We examined the subcellular localization of endogenous SCAF1 by immunofluorescence in Tet-On PEX5^C11A^ cells under unstressed (Dox−) and peroxisomal import stress (Dox+, 1 μg/ml, 2 days) conditions, using ATP5A1 (the α-subunit of the mitochondrial ATP synthase) as a mitochondrial marker (**Fig. 2.A**). Under basal conditions, SCAF1 displayed predominantly nuclear and diffuse cytoplasmic distribution with minimal co-localization with the mitochondrial ATP5A signal, consistent with its canonical annotation as a nuclear splicing factor (*61*, *67*). Upon doxycycline-induced PEX5^C11A^ expression, SCAF1 localization shifted from nuclear to predominantly mitochondrial (**Fig.2.B**). To confirm this striking mitochondrial translocation, we performed immunoblotting analysis using subcellular fractions isolated using a commercial mitochondrial isolation kit (Thermo Fisher #89874). Our SCAF1 antibody detected three isoforms in the cytosolic fraction, with apparent molecular weights of ∼140 kDa, ∼75 kDa, and ∼35 kDa (**Fig. 2.C**). SCAF1 is predicted to have 16 isofroms [16], of which the 139.3 kDa isoform (1312 aa) is the major nuclear form that regulates RNA Polymerase II elongation and global transcription (*68*, *71*). Notably, the 140 kDa and 35 kDa isoforms were not detected in the mitochondrial fraction, while the 75 kDa isoform (likely a precursor) is processed to a slightly smaller 70 kDa band (potentially a mature form following cleavage of a mitochondrial targeting sequence, MTS) (**Fig. 2.D**). Quantification of immunoblots showed that peroxisomal stress induced a 14-fold increase in the mature mitochondrial form of SCAF1 (70 kDa; **Fig. 2.F**), compared with only a 1.7-fold increase in the cytosolic precursor (75 kDa; **Fig. 2.E**). Thus, we identified a unique SCAF1 isoform that translocates to mitochondria under peroxisomal stress.

### SCAF1 represses mitochondrial translation output under peroxisomal stress without affecting mRNA levels

Unlike the nuclear genome, human mitochondria harbor their own circular genome. The mitochondrial genome encodes 13 protein subunits that are indispensable for the assembly and function of oxidative phosphorylation complexes I, III, IV, and V, alongside the 22 tRNAs and 2 rRNAs (12S and 16S) required for their intra-mitochondrial translation. Expression of these mitochondrially encoded genes is regulated by a dedicated nuclear-encoded transcriptional and post-transcriptional machinery, and disruption at any level, from transcription initiation to RNA turnover to ribosome assembly, can impair OXPHOS capacity and mitochondrial membrane potential. Since SCAF1 is a canonical RNA-binding factor known to regulate transcript fate in the nucleus, its stress-induced redistribution to mitochondria raised the question of whether it similarly influences mitochondrial RNA metabolism.

To determine whether SCAF1 regulates mitochondrially encoded protein levels through a transcriptional or post-transcriptional mechanism, we first measured the mRNA expression of all 13 protein-coding mitochondrial transcripts with both mitochondrial ribosomal RNAs (12S and 16S), 15 mitochondrially encoded RNAs in total, by quantitative RT-PCR in Tet-On PEX5^C11A^ cells following siRNA-mediated knockdown of SCAF1 for 2 days (**Fig 3A**). SCAF1 mRNA itself was significantly reduced in the knockdown condition (*p = 0.03), confirming effective gene silencing. In striking contrast, none of the 15 mitochondrially encoded transcripts showed a statistically significant change between control siRNA and SCAF1 siRNA conditions (all comparisons ns**; supplementary Fig. 3A**). The complete absence of any significant mRNA change across the entire mitochondrial transcriptome indicates that SCAF1 does not detectably alter mitochondrial steady state levels and suggests that any downstream effects on mitochondrially encoded protein levels must operate through a post-transcriptional mechanism.

We next assessed the levels of mtDNA-encoded proteins by immunoblotting. Peroxisomal stress led to significant downregulation of most mtDNA-encoded proteins (**Fig. 3B-3C**), consistent with our previous finding that peroxisomal stress inhibits ribosome biogenesis (*25*). Because *PEX5^C11A^* induction relies on low-dose Dox (1 μg/mL), which might inhibit mitochondrial translation. We then performed another immunoblotting in Dox-treated wild-type HEK293 cells and found that low-dose Dox (1 μg/mL) had no effect on mtDNA-encoded protein levels (**Fig. 3D-3E**), indicating that the observed translational inhibition upon *PEX5^C11A^* induction is not attributable to Dox.

In contrast, *SCAF1* KD markedly upregulated most mtDNA-encoded proteins, particularly those of Complexes I and V (**Fig. 3F-3G**). Collectively, these results support a model in which peroxisomal stress promotes mitochondrial localization of SCAF1, which in turn inhibits mitochondrial translation to preserve protein homeostasis and sustain mitochondrial bioenergetic function.

### The N-terminal region of SCAF1 is sufficient for mitochondrial translational repression

SCAF1 is an 11-exon gene encoding a 1,312 amino acid protein organized into three functionally distinct regions: an N-terminal serine/arginine-rich (SR) domain that mediates protein–protein interactions and undergoes phosphorylation-dependent nucleocytoplasmic shuttling, a central RS domain that serves as both a nuclear localization signal and an RNA-binding platform, and a C-terminal CTD-interaction domain (CID) through which SCAF1 engages RNA polymerase II during co-transcriptional splicing (*68*). To establish whether translational repressive function of SCAF1 is autonomous to the mitochondrial compartment and separable from its canonical nuclear splicing activity, we first generated a constitutive SCAF1 knockout stable cell line by CRISPR-Cas9-mediated targeting of exon 6 in HAP1 human near-haploid cells (*92*). The near-haploid nature of HAP1 cells ensures that a single allelic disruption produces a complete null without requiring biallelic editing. Following validation by Sanger sequencing and Western blotting in two independently derived clones, SCAF1 KO cells exhibited markedly elevated MT-ATP6 and MT-ATP8 protein levels compared to wild-type (WT) controls (Fig. **4E-G**), successfully recapitulating earlier siRNA knockdown phenotypes.

**Fig 4.**
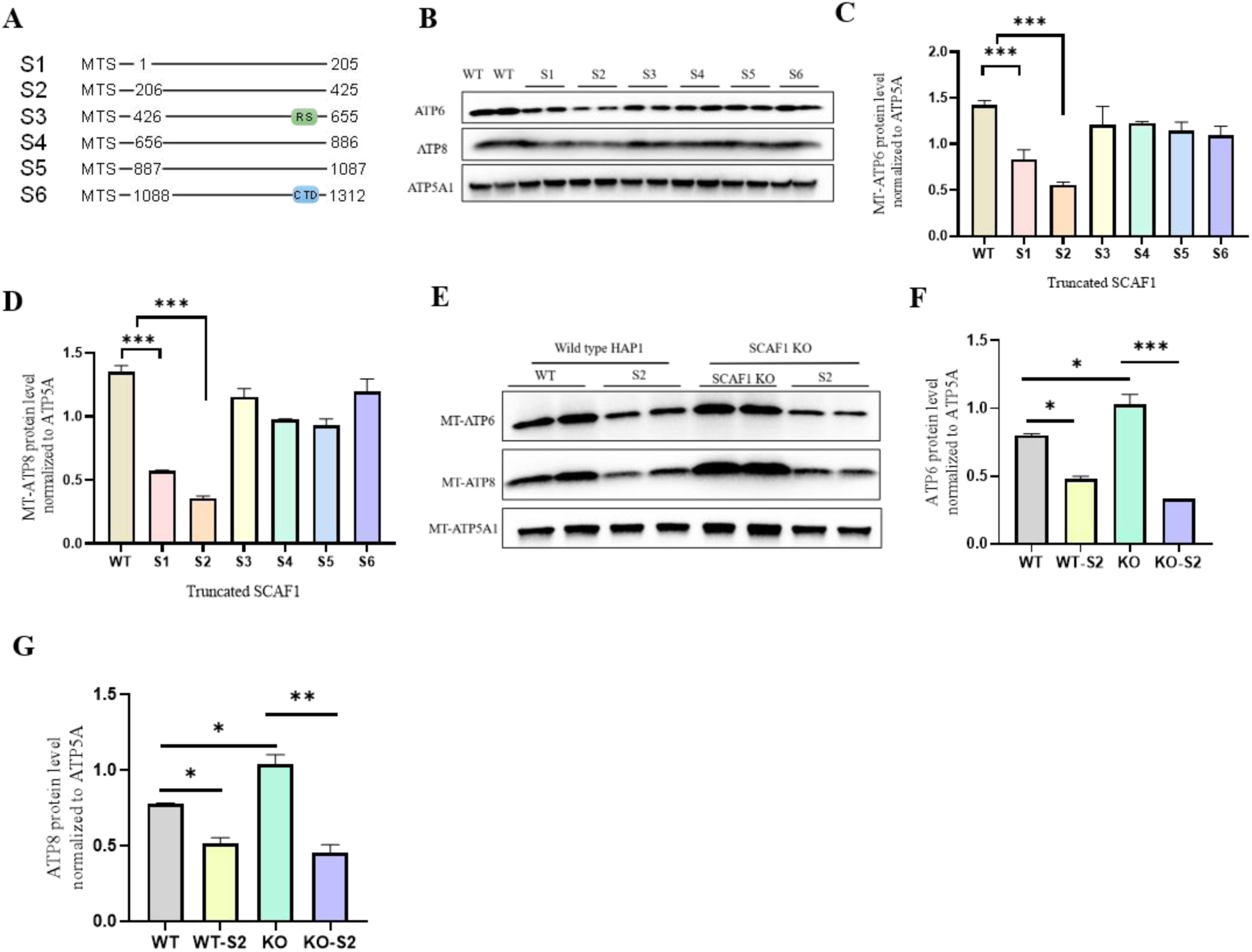
N-terminal region of SCAF1 inhibits mitochondrial translation. (A) Schematic illustration of six truncated SCAF1 constructs. (B) Immunoblot analysis of mitochondrial MT-ATP8 and MT-ATP6 expression upon transient transfection of SCAF1 fragments in WT HAP1 cells (N=2) (C-D) Densitometric analysis for MT-ATP6 and MT-ATP8 respectively normalized to ATP5A (E) Immunoblot analysis of mitochondria MT-ATP8 and MT-ATP6 expression upon transient transfection of fragment S2 in SCAF1 K0 HAP1 cells ( F-G) Densitometric analysis of MT-ATP6 and MT-ATP8 normalized to ATP5A respectiove. N = 2; Student’s t-test. *, *p*<0.05. **, *p*<0.01. ***, *p*<0.001.

To systematically delineate the specific functional domains responsible for this inhibitory activity, we mapped the minimal functional domain by generating a series of six non-overlapping, truncated SCAF1 constructs (S1 to S6) spanning the full sequence. Each fragment was fused to an N-terminal COX8 mitochondrial targeting sequence (MTS) to force mitochondrial localization (**Fig. 4.A**). We transiently expressed these MTS-tagged fragments in WT HAP1 cells and assessed mitochondrially encoded protein levels. Immunoblotting and subsequent densitometric analysis revealed that the N-terminal constructs S1 (amino acids 1 to 205) and, most profoundly, S2 (amino acids 206 to 425) significantly reduced the levels of the mtDNA-encoded proteins MT-ATP6 and MT-ATP8 (Fig. 4, B to D). In contrast, all C-terminal fragments (S3 to S6) including the structured CTD-binding domain previously implicated in RNA polymerase II interactions failed to repress mitochondrial translation.

To confirm the S2 fragment as the minimal effector module, we overexpressed the S2 construct in both WT and SCAF1 KO cells (**Fig. 4.E**). Expression of the S2 fragment in SCAF1 KO cells effectively attenuated the knockout-induced elevation of MT-ATP6 and MT-ATP8, rescuing translational derepression independently of SCAF1’s nuclear activity (**Fig. 4, F and G**). Furthermore, expression of the S2 construct in WT cells drove MT-ATP6 and MT-ATP8 below basal levels in a gain-of-function manner. The nuclear-encoded mitochondrial protein ATP5A1 remained unchanged across all conditions, confirming the specificity of this regulatory mechanism. Together, these data identify the 206- to 425–amino acid region of SCAF1 as the minimal, cell-autonomous mitochondrial effector module, demonstrating that the structured C-terminal domain required for canonical nuclear splicing is entirely dispensable for mitochondrial translational repression.

### The SCAF1 interactome is centered on the mitoribosomal translation machinery

To elucidate the broader molecular functions of SCAF1, we identified its interacting protein network using affinity purification coupled with mass spectrometry (AP-MS). Functional categorization of the resulting SCAF1 interactome demonstrated a significant enrichment of proteins localized to the mitochondria (21%) and those involved in ribosomal functions (13%) (**Fig. 5A**). A detailed analysis of the top-ranked SCAF1-interacting candidates revealed a robust network of proteins critical for mitoribosome assembly—including WBSCR16, DDX28, DHX30, GTPBP10, FASTKD2, FASTKD4, and MALSU1—alongside essential mitochondrial mRNA processing factors (LRPPRC) and chaperones (HSPA9, HSPD1) (**Fig. 5B**).The exclusive matrix localization of these interactors confirms that the processed SCAF1 fragment is successfully imported into the mitochondria, where it physically integrates with the mitoribosomal assembly and folding machinery. By engaging multiple translation-associated factors and chaperones rather than a single target, SCAF1 establishes a broad interaction network that provides the necessary molecular support to maintain mitochondrial translational homeostasis. To independently validate these physical associations and map the specific binding interfaces, we engineered a series of FLAG-tagged SCAF1 truncation mutants equipped with a mitochondrial targeting sequence (MTS): S1 (residues 1–205), S2 (residues 206–425), S5 (residues 887–1087), and S6 (residues 1088–1312). Following FLAG-pulldown, immunoblot analysis of the eluted fractions confirmed that SCAF1 co-immunoprecipitates with a wide array of endogenous mitoribosome assembly factors (**Fig. 5C**).

**Fig. 5.**
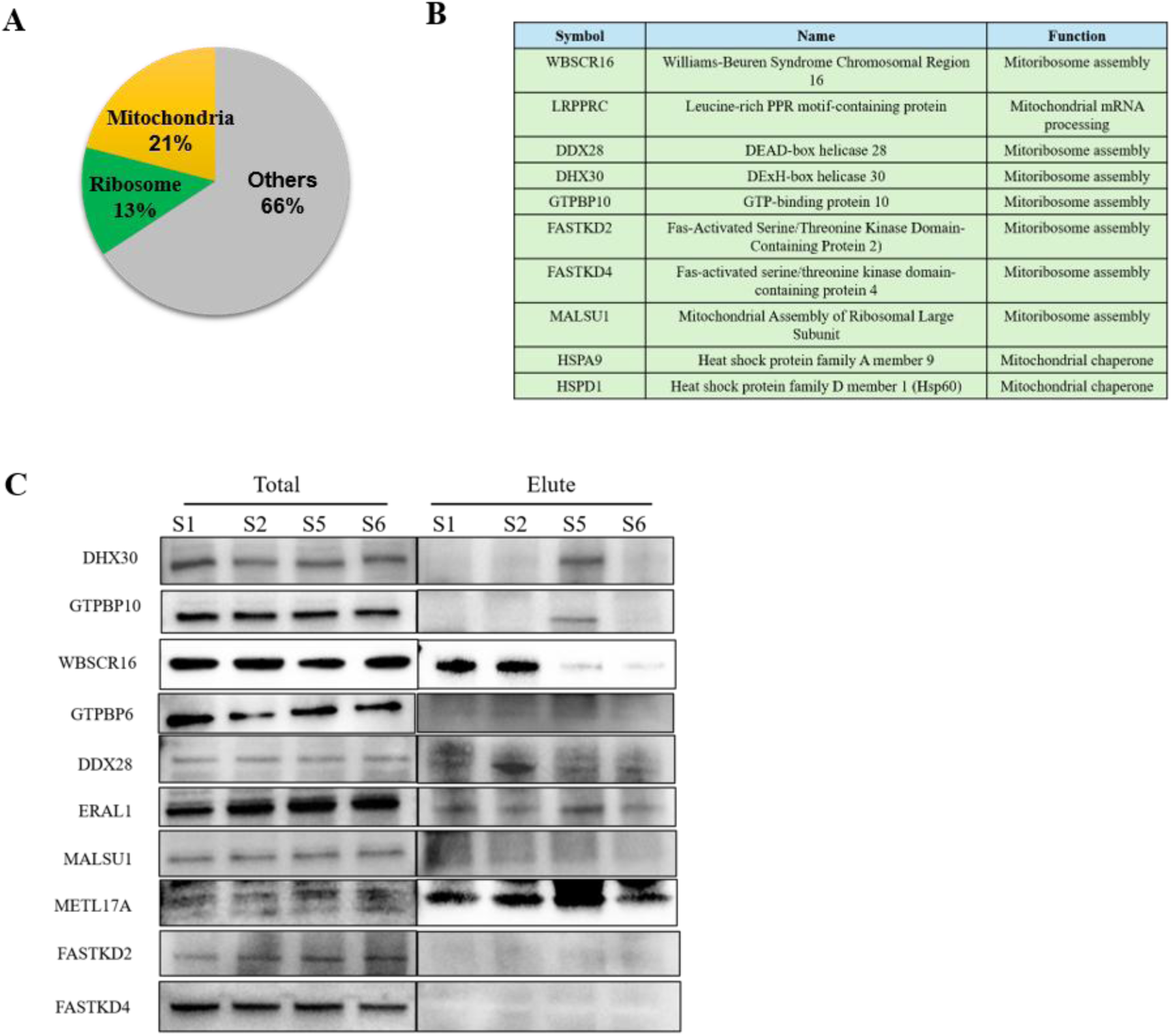
SCAF1 physically associates with mitochondrial and mitoribosome-related proteins. (A) Functional categorization of proteins identified by affinity purification-mass spectrometry (AP-MS) of SCAF1, demonstrating a significant enrichment of mitochondrial (21%) and ribosomal (13%) proteins. (B) Representative list of the top SCAF1-interacting proteins identified by mass spectrometry, highlighting their distinct cellular functions in mitoribosome assembly, mitochondrial mRNA processing, and chaperone activity. (C) Immunoblot analysis of FLAG-pulldowns from cells expressing various FLAG-tagged SCAF1 truncation mutants equipped with a mitochondrial targeting sequence (MTS). The fragments used are S1 (residues 1–205), S2 (residues 206–425), S5 (residues 887–1087), and S6 (residues 1088–1312). Total lysates and eluted fractions were immunoblotted for the indicated mitoribosome assembly factors to map the primary SCAF1 interaction domains.

Integrating these binding profiles with our functional mapping reveals how SCAF1 coordinates mitochondrial translation. While multiple mitoribosome assembly factors anchor broadly across the S1 and S5 regions, the S2 domain (amino acids 206–425) emerges as a critical functional module. Specifically, interaction analyses highlight DDX28 and WBSCR16 as key candidates interacting within this window. Because our preceding domain analysis demonstrated that the S2 fragment is uniquely both necessary and sufficient to drive the repression of mtDNA-encoded proteins (such as MT-ATP6 and MT-ATP8), these targeted interactions suggest that SCAF1 leverages specific sub-domains to anchor to the mitoribosome machinery while utilizing the S2 module to directly execute translational control.

### SCAF1 controls the mitoribosomal subunit joining step to regulate 55S monosome production

The interaction of SCAF1 with key mitoribosome assembly factors (**Fig. 5B**) prompted us to test whether SCAF1 regulates mitochondrial translation by modulating mitoribosome biogenesis. Assembly of the mitoribosome is a tightly regulated process requiring incorporation of nuclear-encoded proteins and mitochondrial rRNAs into functional small and large subunits (*93*, *94*).

To investigate the potential role of SCAF1 in mitoribosome assembly, we performed sucrose density gradient centrifugation, a powerful biochemical technique that separates mitochondrial ribosomal components based on their size, shape, and density in a linear 10-30% sucrose gradient during ultracentrifugation (**Fig. 6A**) (*95*). This approach resolves assembly intermediates, the 28S mitoribosome small subunit (mtSSU), the 39S mitoribosome large subunit (mtLSU), and the fully assembled 55S monosome. Using the antibodies against mtLSU and mtSSU markers (MRPL11 and MRPS15 respectively), we profiled the mitoribosome assembly in a SCAF1 stable knockout (KO) cell line that we generated via CRISPR-Cas9 technology in human near-haploid HAP1 cells (**Fig. 6B**). Sequencing analysis confirmed a deletion of 19 nucleotides (190–195) in SCAF1, resulting in a frameshift mutation at amino acid 136 (**Fig. 6B**). Immunoblot analysis verified that the KO is a null mutant with no detectable expression of the 70 kDa SCAF1 isoform in isolated mitochondria (**Fig. 6C).**

**Fig 6.**
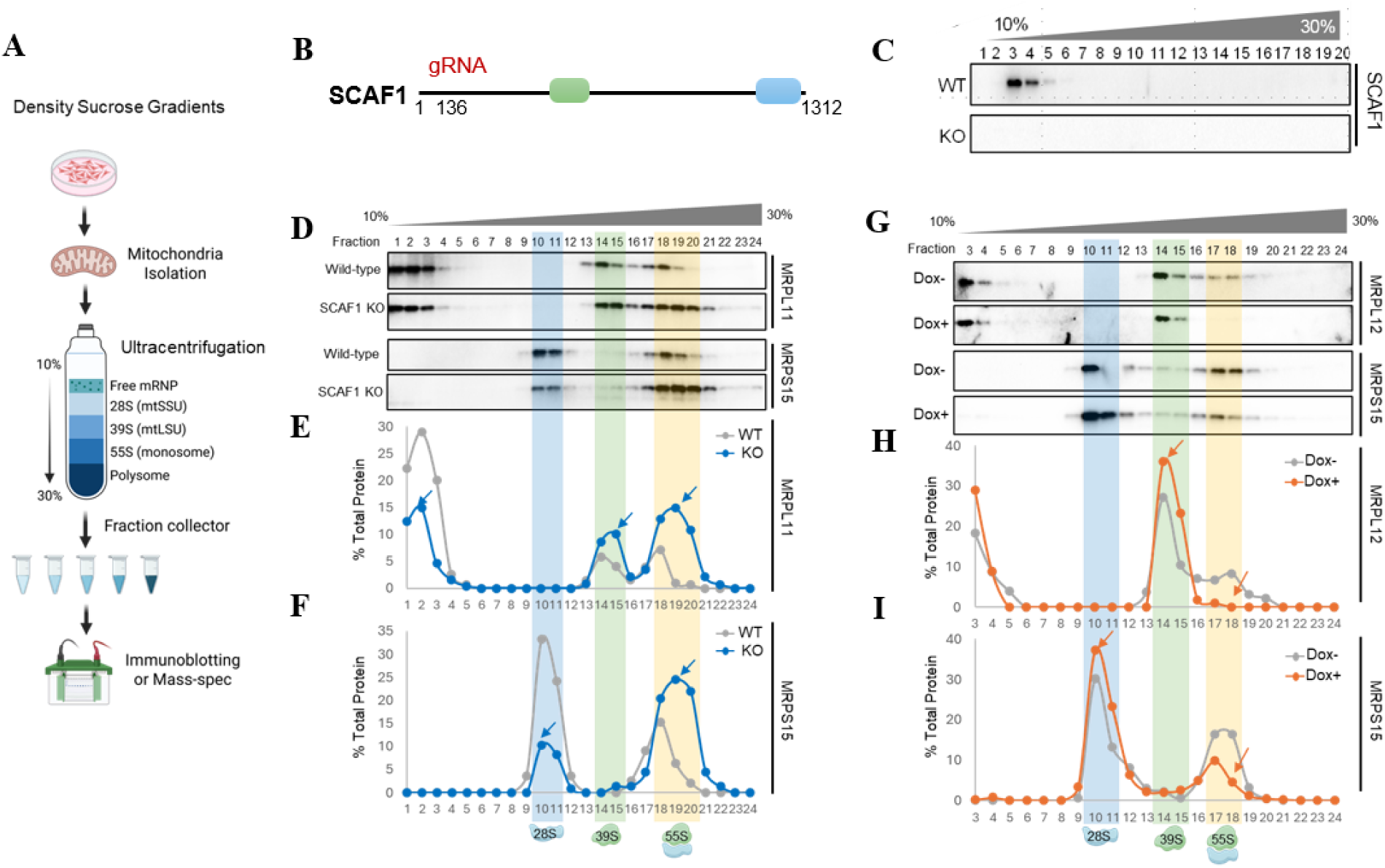
SCAF1 negatively regulates mitoribosome assembly. (A) Schematic of the sucrose density gradient centrifugation workflow used for mitoribosome profiling. (B) Diagram showing the CRISPR gRNA target site used to generate the SCAF1 KO. (C) Immunoblot analysis of isolated mitochondrial fractions confirming that the *SCAF1* KO is a null mutant. (D, G) Immunoblotting of mitochondrial fractions separated by 10-30% sucrose density gradient centrifugation from *SCAF1* KO cells or *PEX5^C11A^*-expressing HEK293 cells. MRPL11 or MRPL12 antibodies were used to detect the 39S mtLSU, and MRPS15 as a marker for the 28S mtSSU. (E, F, H, I) Quantification of mtLSU, mtSSU, and monosome, expressed as the percentage of total band intensity across gradient fractions.

In alignment with the increased mitochondrial translation in *SCAF1* KD cells (**Fig. 6F-6G**), *SCAF1* KO led to a marked increase in mature 55S monosome and 39S mtLSU, accompanied by reduced levels of 28S mtSSU and pre-mtLSU (lighter fraction, #1-4) (**Fig. 6D-6F**). Immunoblots with anti-MRPL11 revealed three distinct peaks corresponding to pre-mtLSU, mature mtLSU (39S), and monosome (55S) (**Fig. 6E**), while anti-MRPS15 showed two peaks for mtSSU (28S) and monosome (55S) (**Fig. 6F**). This shift toward mature monosomes in *SCAF1* KO cells indicates that SCAF1 acts as a negative regulator of mitoribosome assembly, particularly at the subunit joining step leading to monosome formation.

In contrast, sucrose density gradient analysis of *PEX5^C11A^*-expressing cells (Dox+) revealed an opposite pattern (**Fig. 6G-6I**): a significant reduction in 55S monosomes with concomitant accumulation of 39S mtLSU (**Fig. 6H**) and 28S mtSSU (**Fig. 6I**). This impairment in monosome formation likely underlies the previously observed downregulation of mtDNA-encoded protein expression under peroxisomal stress (**Fig. 2B-2C**).

### SCAF1 is recruited to mitochondrial RNA granules during peroxisome stress to regulate early assembly checkpoints

To determine the spatial dynamics of SCAF1 and its interacting partners during mitoribosome biogenesis, we performed sucrose density gradient fractionations on mitochondria isolated from wild-type (WT) and SCAF1 knockout (KO) cells (**Fig. 7, A-E**). In WT mitochondria, SCAF1 co-sediments with the assembly factor WBSCR16 and RNA processing factors FASTKD4 and LRPPRC predominantly in the lighter gradient fractions (fractions 1–6), corresponding to mitochondrial RNA granules and free ribonucleoproteins (mRNPs). Because the 28S and 39S mitoribosomal subunits are formed independently rather than sequentially, these early granule stages represent a critical regulatory node prior to subunit maturation. In SCAF1 KO mitochondria, the peak distributions of WBSCR16, FASTKD4, and LRPPRC exhibited noticeable architectural shifts within these early fractions (**Fig. 7, C-E**). This indicates that basal SCAF1 is required to maintain the native stoichiometric organization of these pre-assembly complexes.

**Fig 7:**
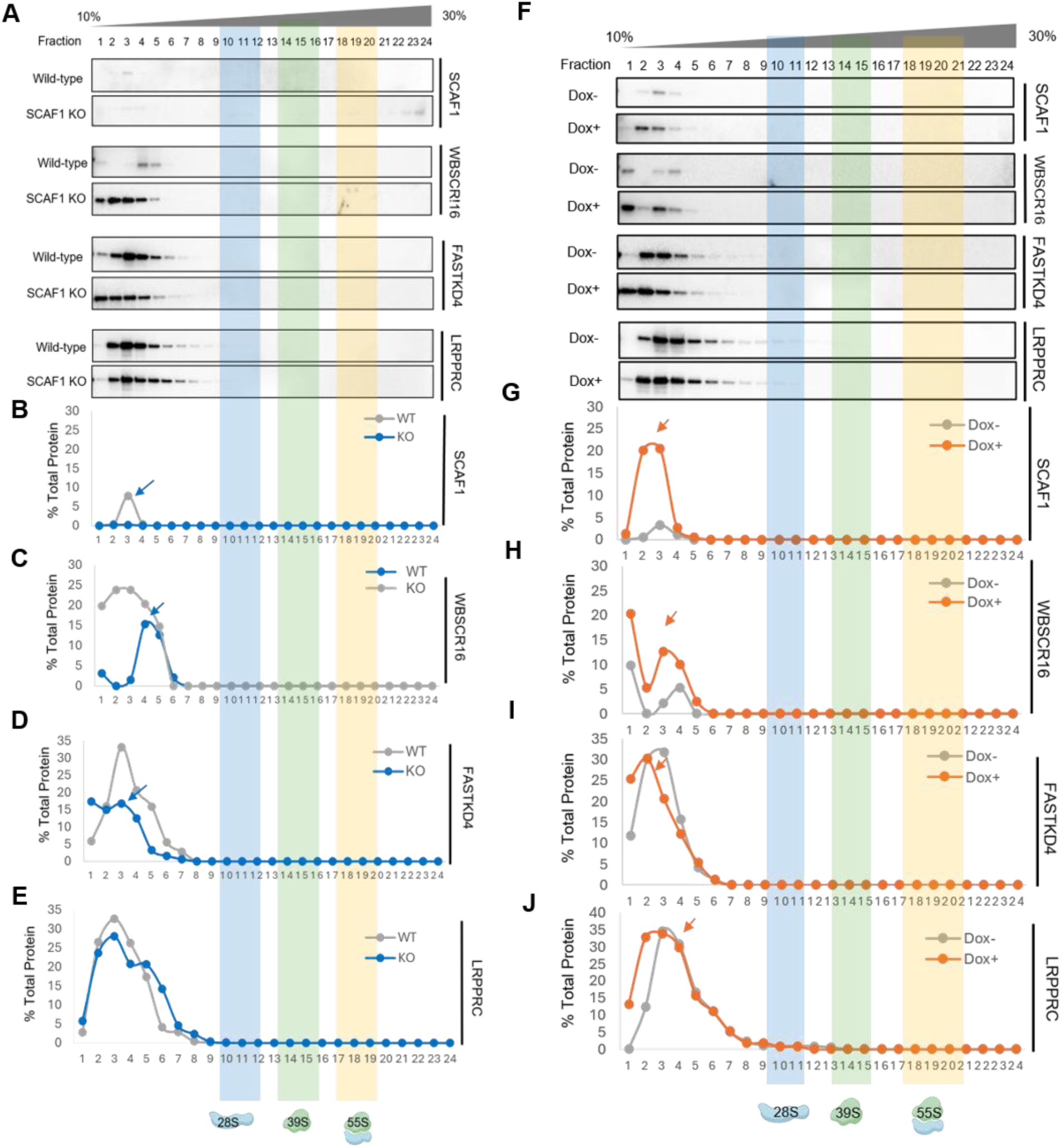
SCAF1 is recruited to RNA granule under peroxisome stress and regulates assembly checkpoints. (A-E) Western blots and quantification (% total protein per fraction) For SCAF1, WBSCR16, FASTKD4 and LRPPRC across 24 fractions from WT and SCAF1 KO mitochondria. Blue arrows indicate shifts in the peak for each antibody tested. Colored zones: blue is 28S, green is 39 S yellow is polysome. (F-J) Westen blots and quantification for SCAF1, WBSCR16, FASTKD4, and LRPPRC across 24 fractions from Dox- and Dox + mitochondria. Orange arrows indicate the shifts in the peaks along the profile.

We next examined how this network responds to peroxisome stress. Following doxycycline-induced peroxisome dysfunction (Dox+), gradient profiling revealed a striking spatial redistribution. SCAF1 was robustly recruited to, and highly enriched within, the early RNA granule fractions (**Fig. 7, F-G**). Concurrently, WBSCR16, FASTKD4, and LRPPRC displayed pronounced, coordinated shifts, accumulating prominently alongside SCAF1 in these lighter molecular weight complexes (**Fig. 7, H-J**). Crucially, despite this accumulation within RNA granules, our earlier transcriptomic analysis confirmed that steady-state mitochondrial mRNA levels remain entirely unperturbed. This demonstrates that SCAF1 does not induce transcript degradation but instead acts strictly as a localized translational brake.

Integrating these spatial dynamics with our structural interaction mapping reveals a comprehensive mechanism for peroxisome-to-mitochondria stress signaling. SCAF1 utilizes its S1 and S5 domains to broadly dock onto early mRNP complexes. Once physically anchored within the granule under stress conditions, SCAF1 deploys its minimal functional effector module, the S2 domain (amino acids 206–425) to specifically engage critical assembly factors, most notably WBSCR16 and DDX28. This targeted, S2-mediated interaction effectively stalls the independent maturation pathways of the mitoribosomal subunits at the foundational RNA granule stage, preventing their downstream joining into functional 55S monosomes. By imposing this early assembly checkpoint, SCAF1 directly represses the translation of key mtDNA-encoded components (e.g., MT-ATP6 and MT-ATP8) and dampens oxidative phosphorylation, allowing the cell to rapidly adapt to peroxisomal dysfunction while preserving its intact mitochondrial transcriptome.

Together, these findings demonstrate that peroxisome stress recruits SCAF1 to mitochondrial RNA processing centers and dynamically alters mitoribosomal assembly checkpoints to regulate translational homeostasis.

### Peroxisomal dysfunction and mitochondrial SCAF1 accumulation drive cellular senescence in human fibroblasts

To determine whether peroxisomal dysfunction directly impacts cellular longevity and fates, we evaluated markers of cellular senescence in human IMR90 diploid fibroblasts following peroxisome disruption. Knockdown of the essential peroxisomal import receptor PEX5 via small interfering RNA (siRNA) significantly increased senescence-associated β-galactosidase (SA-β-gal) activity compared to scramble siRNA controls (**Fig. 8, A-B**). Quantitative analysis revealed that PEX5 depletion led to a nearly sixfold increase in SA-β-gal–positive cells (∼38% versus ∼6% in scramble controls; Fig. 8B). To establish whether this senescence phenotype was accompanied by canonical molecular pathways, we analyzed the expression of key cell cycle regulators and components of the senescence-associated secretory phenotype (SASP) by quantitative reverse transcription polymerase chain reaction (RT-qPCR). PEX5 knockdown induced a robust transcriptional upregulation of the cyclin-dependent kinase inhibitor *CDKN1A* (p21; ∼3-fold), alongside prominent SASP proinflammatory cytokines and chemokines, including *CXCL2* (∼2.5-fold), *IL6* (∼2-fold), and *IL7* (∼2-fold) (**Fig.8.E**). Together, these findings demonstrate that peroxisomal impairment drives primary human fibroblasts into a state of irreversible growth arrest accompanied by a distinct secretory phenotype.

**Fig 8.**
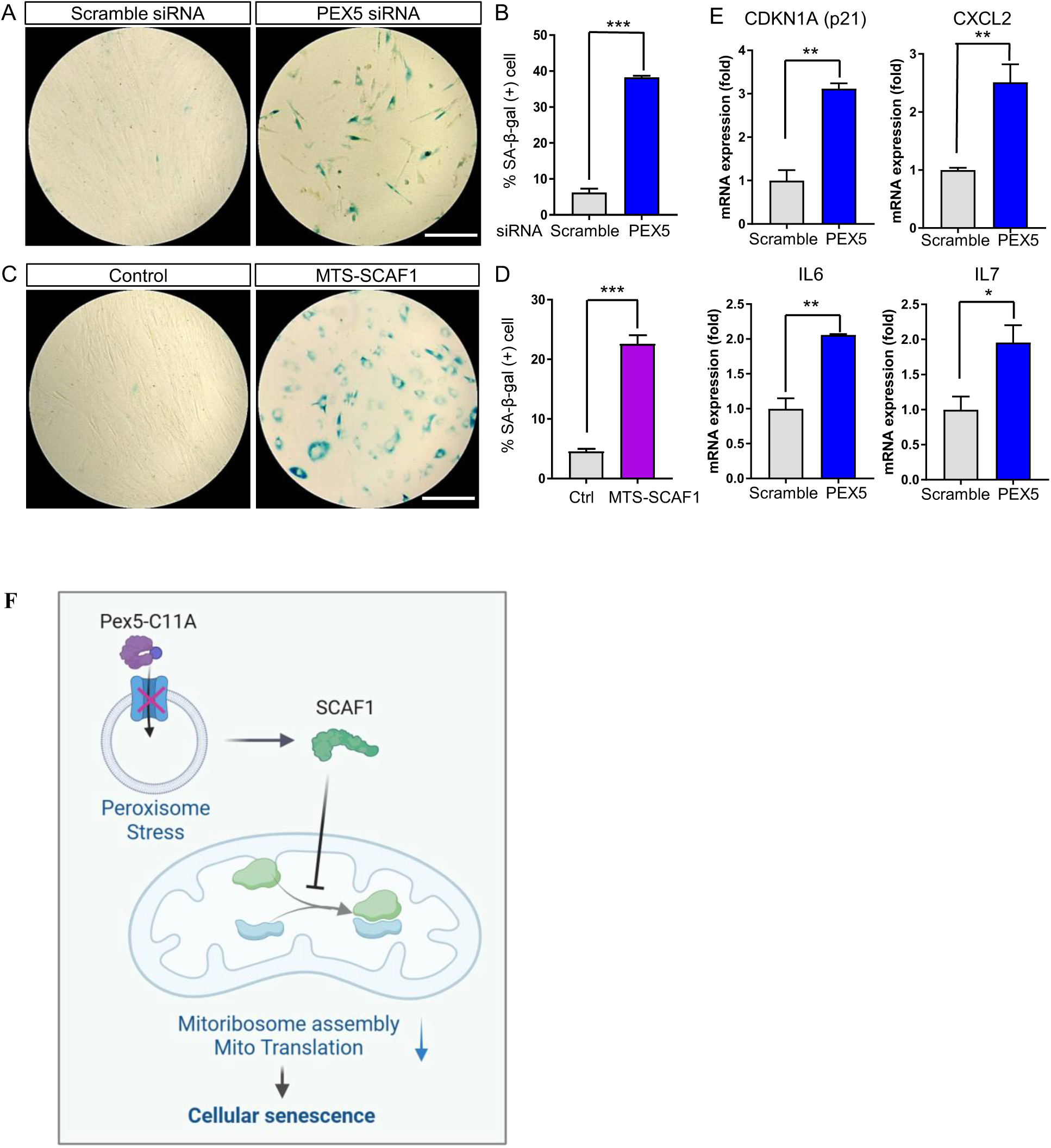
Peroxisomal dysfunction promotes cellular senescence in human fibroblasts. (A, C) Senescence-associated β-galactosidase (SA-β-gal) staining of human IMR90 fibroblasts transfected with *PEX5* siRNA or MTS-SCAF1 constructs. Scale bar, 260 μm. (B, D) Quantification of SA-β-gal-positive cells under *PEX5* KD or SCAF1 overexpression. N=3, Student’s t-test. (E) Quantitative RT-PCR analysis of senescence-associated gene expression (p21, CXCL2, IL-6, and IL-7) in IMR90 cells transfected with scramble or *PEX5* siRNA. N=3, Student’s t-test. (F)The conceptual framework for SCAF1-mediated peroxisome-mitochondria crosstalk drives cellular senescence.

Because peroxisomal stress promotes the mitochondrial translocation and accumulation of SCAF1 to regulate translational homeostasis, we next asked whether forced mitochondrial targeting of SCAF1 is sufficient to recapitulate the senescent phenotype observed during peroxisomal failure. We overexpressed an N-terminally tagged mitochondrial targeting sequence construct of SCAF1 (MTS-SCAF1) or an empty vector control in human IMR90 fibroblasts (). Expressing MTS-SCAF1 alone was sufficient to induce robust SA-β-gal staining, driving a more than fivefold increase in the proportion of senescent cells relative to controls (∼22% versus ∼4%; **Fig. 8, C-D**). These results indicate that forced mitochondrial recruitment of SCAF1 acts as a downstream effector of peroxisomal stress to autonomously induce cellular senescence in human cells.

## DISCUSSION

Peroxisomes and mitochondria are metabolically interdependent organelles that cooperate in essential pathways, including fatty acid *β*-oxidation, reactive oxygen species (ROS) homeostasis, and ether lipid synthesis. While functional decline in either organelle is known to trigger systemic metabolic failure, the precise molecular mechanisms by which cells sense peroxisomal dysfunction and coordinate retro-signaling responses to mitochondrial targets have remained largely obscure. Here we identify SCAF1 as a previously unrecognized molecular relay that couples peroxisomal import failure to mitochondrial translational repression and protective cellular senescence. In contrast to earlier work that catalogued secondary mitochondrial defects downstream of peroxisomal dysfunction, our data reveal an active, adaptive signaling axis in which a canonical nuclear splicing factor is selectively processed and redirected to mitochondria, where it imposes an early checkpoint on mitoribosome assembly. This mechanism prioritizes mitochondrial proteostasis over maximal bioenergetic output and ultimately drives a senescence program that we interpret as protective in the context of aging and chronic organelle stress.

The peroxisome–mitochondria axis has been defined by metabolite exchange, shared fission machinery, and membrane contact sites (*14*, *30*, *30*, *96–98*). Peroxisomal β-oxidation generates acetyl-CoA and shortened acyl chains that feed directly into the mitochondrial TCA cycle and OXPHOS (*14*, *96*), while fission GTPase DRP1 and its receptor MFF are dually targeted to both organelles, physically coupling their division cycles (*97*, *98*). Membrane contact sites mediated by ACBD5 and VAPB, and by MIRO1, tether peroxisomes to the ER and to mitochondria respectively, enabling direct metabolite and lipid transfer (*30*, *99*). Classic studies established that peroxisomal matrix-import defects, as occur in Zellweger spectrum disorders caused by pathogenic *PEX* gene variants, produce secondary mitochondrial abnormalities including impaired OXPHOS, cristae structural defects, and elevated mitochondrial superoxide (*26*, *100*). Yet the molecular sensors and effectors that detect peroxisomal failure and actively reprogram mitochondrial gene expression remained undefined. Prior work on peroxisome–mitochondria crosstalk has focused primarily on metabolite flux, shared fission GTPases including DRP1 and MFF, and membrane contact sites (*30*, *97*). Our CRISPR screen and subsequent mechanistic dissection uncover a distinct layer of regulation: a stress-induced isoform of SCAF1 that relocates to mitochondria and directly engages the mitoribosomal assembly machinery. This represents, to our knowledge, the first demonstration that a nuclear SR-family splicing factor can function as a peroxisome-to-mitochondria stress transducer. The finding expands the emerging concept that nuclear RNA-binding proteins can acquire organelle-specific roles under stress and places SCAF1 within a growing list of dual-compartment regulators that coordinate nuclear and mitochondrial responses to metabolic perturbation (*101*, *102*).

The mobilization of SCAF1 raises fundamental questions regarding the upstream signals that govern its proteolytic processing and mitochondrial translocation. Under homeostatic conditions, SCAF1 is confined to the nucleus. However, peroxisomal dysfunction rapidly elevates cytosolic reactive oxygen species (ROS) and disrupts intracellular redox poise,(*103–106*) while simultaneously driving the accumulation of unoxidized very-long-chain fatty acids (VLCFAs) (*14*, *107*). We posit that the stress-induced oxidation of conserved cysteine residues within the N-terminal SR domain of SCAF1, or lipid-mediated alterations to its conformation, unmasks a cryptic mitochondrial targeting sequence (MTS). This would parallel the redox-regulated translocation kinetics observed for cytosolic factors like Nrf2 and p53 (*102*, *108*). Furthermore, the robust activation of both the integrated stress response (ISR) and the mitochondrial unfolded protein response (UPR^mt^) observed in our transcriptomic data suggests that stress-dependent alternative splicing may synergistically favor the expression of an import-competent SCAF1 isoform (*81*, *85*). Defining the precise cytosolic or mitochondrial proteases responsible for SCAF1 maturation will be a critical next step in deciphering this signaling relay.

Our domain mapping and interactome data indicate that the S2 region (residues 206–425) is the minimal effector module for translational repression, while broader domains (S1 and S5) help anchor SCAF1 to early mRNP complexes. We speculate that S2 contacts a specific surface on WBSCR16 or DDX28 that is required for subunit joining, thereby sterically or allosterically hindering progression from free 28S/39S particles to mature 55S monosomes. This model is consistent with the gradient shifts we observe and with known roles of these factors in early assembly checkpoints (*47*, *50*, *51*, *109*, *110*). Whether SCAF1 also modulates RNA helicase activity, rRNA modification, or chaperone recruitment remains an open question. Structural studies of the SCAF1–WBSCR16/DDX28 interface, or proximity-labeling approaches under stress versus basal conditions, would test this assumption directly. Notably, no prior report has described SCAF1 (or any closely related SR protein) binding the mitoribosome or regulating mitochondrial translation in any model system, underscoring the novelty of the interaction network we define.

Mitoribosome biogenesis is a spatially organized, multi-step process that begins in the RNA granule, a membraneless compartment adjacent to the mitochondrial nucleoid where mt-rRNA processing, early ribosomal protein assembly, and rRNA modification are coordinated.(*38*, *41*, *50*)Sucrose density gradient sedimentation provides high-resolution evidence that SCAF1 acts at the granule stage rather than at downstream assembly steps: under peroxisomal stress, SCAF1 is confined to low-density fractions 1–5 corresponding to granule-associated species, and is absent from 39S, 55S monosome, and polysome fractions. This mechanistic interpretation is strongly supported by published mouse knockout models: loss of TFB1M, which dimethylates the 3′-loop of 12S rRNA in the small subunit, causes complete loss of assembled 28S subunits with concomitant large subunit accumulation, demonstrating that an upstream rRNA maturation defect propagates asymmetrically to block final monosome formation.(*42*, *111*). Similarly, deletion of MTERF4, which recruits the NSUN4 methyltransferase to late 39S intermediates, stalls assembly factors on a near-mature large subunit that cannot proceed to monosome formation. The published in vivo models demonstrate that upstream rRNA-processing lesions, and our data position SCAF1 as the upstream initiating event in this cascade (*111*).

WBSCR16 behavior in SCAF1 knockout cells provides direct mechanistic definition of this pipeline. In the absence of SCAF1, WBSCR16 advances from granule fractions into 28S-associated fractions, indicating that SCAF1 normally restrains the WBSCR16-mediated granule-to-ribosome transition.(*109*, *110*) The invariant distribution of LRPPRC across all conditions confirms that SCAF1 acts specifically on the rRNA processing branch of granule biology, leaving the LRPPRC– SLIRP mRNA stability axis fully intact.(*52*, *53*) This specificity carries a profound functional consequence: by suppressing ribosome supply rather than mRNA availability or ribosome activity, SCAF1 achieves simultaneous, stoichiometrically balanced suppression of all 13 mitochondrially encoded OXPHOS subunits while preserving the mRNA template pool for rapid translational reactivation once peroxisomal homeostasis is restored.

Several proteins share partial phenotypic overlap with SCAF1 yet operate through distinct mechanisms. LRPPRC and SLIRP control mitochondrial mRNA stability and polyadenylation (*53*, *112*); their loss reduces rather than elevates mtDNA-encoded protein levels. FASTKD-family members and GRSF1 act within RNA granules to promote, rather than restrain, transcript maturation (*56*, *57*, *113*, *114*). On the nuclear side, other SR proteins can translocate under stress, but none have been shown to impose a selective block on mitoribosome assembly. The closest functional parallels may be stress-induced translational repressors such as the ISR kinase PERK or the mitochondrial protease Lon, both of which limit protein synthesis to preserve proteostasis (*115– 118*). SCAF1 is unique, however, in coupling an upstream peroxisomal cue to a post-transcriptional mitochondrial checkpoint via isoform-specific translocation. This dual-compartment logic has not been reported for any other splicing factor or mitoribosome interactor.

The logic of targeting the RNA granule rather than downstream assembly steps reflects a principle of energetic economy. Ribosome biogenesis is the most resource-intensive biosynthetic process in the cell, consuming the majority of cellular transcriptional and translational capacity at steady state.(*119–121*) Once a 39S assembly intermediate exits the granule and enters the matrix phase of biogenesis, a substantial investment of ribosomal proteins, assembly GTPases, rRNA modification enzymes, and ATP has already been irrevocably committed. Blocking assembly at the granule—before this commitment—represents the earliest, most energetically favorable, and most strategically positioned intervention point in the entire pathway. This consideration is particularly acute under peroxisomal dysfunction, where substrate supply to the mitochondrial TCA cycle and OXPHOS is already compromised by disrupted peroxisomal β-oxidation; continued assembly of OXPHOS machinery without adequate metabolic substrate would represent an energetically catastrophic mismatch.(*14*, *96*, *122*) Throttling ribosome supply at the granule thus provides proportional, substrate-matched, and rapidly reversible control over OXPHOS capacity, conceptually analogous to the TOR-mediated regulation of cytosolic ribosome biogenesis in response to nutrient availability, in which ribosome production—not ribosome activity—is the primary adjusted variable.(*123–125*)

The disease relevance of SCAF1 is supported by multiple converging lines of evidence. SCAF1 overexpression in aggressive ovarian and breast cancers (*67*, *68*) may reflect constitutive mitoribosome suppression that sustains the Warburg phenotype. SCAF1 loss as a tumour suppressor in pancreatic ductal adenocarcinoma (*70*) would remove this brake, potentially driving OXPHOS overactivation. In Zellweger spectrum disorders, the chronic peroxisomal dysfunction caused by *PEX* mutations directly recapitulates the PEX5^C11A^ model used here (*14*, *17*, *18*, *26*); the secondary mitochondrial pathology in these patients (*26*) may be partly explained by constitutive SCAF1 deployment creating a permanent translational brake that cannot resolve because the upstream peroxisomal insult persists. The genetic association of SCAF1 with cardiac conduction phenotype (*72*) is intriguing given the dependence of cardiac tissue on mitochondrial OXPHOS. Whether SCAF1 variants alter the responsiveness or kinetics of the SCAF1-mediated translational brake, thereby perturbing cardiac energetics under metabolic stress, represents a testable hypothesis for future genetics and physiological studies.

The most immediate disease context is the Zellweger spectrum of peroxisome biogenesis disorders (ZSD), a group of autosomal recessive conditions caused by mutations in *PEX* genes that abolish peroxisomal matrix protein import and produce both non-functional peroxisomes and secondary mitochondrial dysfunction contributing to progressive neurodegeneration, metabolic crisis, and early mortality (*15*, *26*, *126*). Mitochondrial structural abnormalities—including cristae disorganization, impaired respiratory chain activity, and morphological fragmentation—are directly caused by defective peroxisomal biogenesis in mouse models of ZSD and in patient fibroblasts, establishing that these are not incidental features but mechanistically linked consequences of peroxin loss (*26*, *126*, *127*). Our finding that SCAF1 drives a protective senescence program raises the possibility that this pathway is activated in ZSD patient cells as an adaptive attempt to limit further organelle damage; however, whether SCAF1 levels or mitochondrial localization are altered in Zellweger fibroblasts or animal models has not been examined and warrants direct investigation. More broadly, peroxisomal function declines progressively with chronological age across multiple tissues (*13*, *128–131*). Catalase activity falls by 30–40% in aged rodent liver, peroxisomal matrix protein import efficiency is compromised in aged human fibroblasts, and fatty acid β-oxidation capacity deteriorates with age—collectively producing lipid accumulation, redox imbalance, and secondary mitochondrial deterioration that are recognized hallmarks of cellular aging (*13*, *128–131*). We therefore propose that chronic, subclinical peroxisomal stress in aged tissues constitutively engages the SCAF1 pathway, initially promoting a protective senescence program that eliminates cells with compromised bioenergetics, but ultimately becoming deleterious when senescent cells accumulate and sustain a pro-inflammatory senescence-associated secretory phenotype (SASP) (*132–138*). This transition from acute protection to chronic pathology is mechanistically supported by recent evidence that mitochondrial dysfunction in senescent cells drives mtDNA release into the cytosol, activating cGAS–STING signaling and amplifying SASP gene expression (*137*) a pathway directly downstream of the mitoribosomal translation suppression that SCAF1 imposes. In this framework, SCAF1 sits at the intersection of two primary aging axes: organelle quality control and the senescence–SASP response(*110*, *132*, *138*). In this framework, SCAF1 sits at the intersection of two major aging axes: organelle quality control and the senescence response. Therapeutic modulation of SCAF1 mitochondrial activity—either to enhance protective senescence under acute peroxisomal stress or to dampen chronic SASP-driven inflammation in aged tissues—could therefore be explored in models of ZSD and age-related metabolic decline.

The broader conceptual significance of these findings is threefold. First, SCAF1 establishes mitoribosome subunit joining as a dynamically regulated node subject to extrinsic stress control, analogous to the well characterized eukaryotic translation initiation checkpoints such as elF2α phosphorylation in the ISR, (*84*, *85*) but operating at the level of ribosome biogenesis rather than ribosome activity. Second, SCAF1 defines a new class of inter-organelle relay in which stress signals originating in one organelle are transduced by a nuclear protein that is proteolytically processed distinct from mitochondria-to-nucleus retrograde signaling in the mtUPR (*78*, *81*), the DELE1 -HR1 cytosolic relay, (*87*, *110*) and nucleus to mitochondria anterograde control by nuclear -encoded mitochondrial biogenesis factors (*82*, *83*), and instead representing a tripartite peroxisome-nucleus-mitochondria communication axis. Third, SCAF1 exemplifies the principle that moonlighting proteins acquire regulatory function in stress contexts that are orthogonal to their homoeostatic roles (*74–77*) with the N-terminal intrinsically disordered region serving as a modular, portable effector that can be co-opted for distinct regulatory outputs depending on subcellular localization.

The present study is limited by its reliance on immortalized and near-haploid cell lines for most mechanistic work; validation in primary neurons, glia, and aged tissues will be essential. The precise protease that generates the mature mitochondrial isoform, the full repertoire of upstream signals, and the long-term organismal consequences of SCAF1-mediated translational control remain to be defined. Nevertheless, the data establish SCAF1 as a, stress-responsive node in peroxisome–mitochondria communication and open a new avenue for understanding how cells adapt mitochondrial gene expression when peroxisomal homeostasis fails, particularly in the context of aging.

In summary, SCAF1 exemplifies an adaptive strategy in which a nuclear RNA-processing factor is repurposed to safeguard mitochondrial proteostasis and promote protective senescence under peroxisomal stress. This discovery adds a new molecular layer to organelle crosstalk and highlights SCAF1 as a potential target for interventions aimed at mitigating secondary mitochondrial pathology in peroxisomal disorders and age-related decline. SCAF1 defines a new class of stress-responsive inter-organelle relay: a nuclear RNA-processing factor that is proteolytically converted into a mitochondrial negative regulator of mitoribosome biogenesis when peroxisomal homeostasis is compromised. By limiting ribosome supply rather than ribosome activity, SCAF1 provides proportional, reversible control of the entire mitochondrially encoded proteome, while maintaining OXPHOS stoichiometry during peroxisomal stress. These findings identify the mitoribosome subunit joining checkpoint as a dynamically regulated node subject to extrinsic stress control, establish a new conceptual axis of peroxisome-to-mitochondria communication, and open new directions for understanding how organelle dysfunction propagates across the mitochondrial gene expression network in metabolic disease, neurodegeneration, and ageing.

## MATERIALS AND METHODS

### Plasmid Construction

Human PEX5 cDNA was purchased from Dharmacon Mammalian Gene Collection. The hPEX5 was amplified by PCR using the forward and reverse primers (5′-CACTATAGGGAGACCCAAGCTTATCTAGACATGGCAATGCGGGAGCT-3′ and 5′-TCTTACTTGTCA TCGTCGTCCTTGTAGTCGCCCTGGGGCAGGCC-3′) and introduced between XhoI and BamHI sites in c-Flag pcDNA3 (Addgene #20011) to generate Flag-tagged hPEX5 using NEBuilder HiFi DNA assembly Master mix (New England Biolabs). Site-directed mutagenesis for amino acid substitution (cysteine to alanine at position 11, C11A) was performed using the Q5 Site-directed mutagenesis kit (New England Biolabs) according to the manufacturer’s instruction. The primers for the PEX5^C11A^ mutant were 5′-GGAGGCCGAAgctGGGGGTGCCA ACC-3′ and 5′-ACCAGCTCCCGCATTGCC-3′. To generate tetracycline-inducible PEX5C11A plasmid, we modified pMK243 (Tet-OsTIR1-PURO) from Masato Kanemaki (Addgene #72835). pMK243 was digested by BgIII and MluI to remove the OsTIR sequence. Flag-PEX5^C11A^ was amplified by PCR using the forward and reverse primers (5′-gattatgatcctctagacatatgctgcagattacttgtcatcgtcgtccttgtagt-3 and 5′-tcctaccctcgtaaagaattcgcggccgcaatggcaatgcgggagctggt-3′) and introduced between BgIII and MluI sites in digested pMK243 plasmid to generate TetPEX5^C11A^-PURO plasmid (*25*). All plasmids are confirmed by Sanger sequencing. Additional plasmids to identify the functional part of SCAF1 was ordered from Twist Bio. Details are in the supplementary Table.

### Generation of CRISPR knock-in HEK293 cells expressing PEX5

HEK293 cells were cultured in Dulbecco’s Modified Eagle Medium (DMEM) containing 10% (v/v) fetal bovine serum, with penicillin and streptomycin. Cells were incubated in a 37 °C incubator in an atmosphere of 5% CO2 in air. To generate a stable cell line, we followed the protocol described by Natsume et al. (*139*). 1 × 10^6^ HEK293 cells were plated in one well of a 6-well plate. After 24 h, 800 ng of AAVS1 T2 CRISPR in pX330 (Addgene #72833) and 1 μg of Tet-PEX5^C11A^-PURO were transfected using Effectene (Qiagen) according to the manufacturer’s instructions. After 48 h, the cells were detached and diluted at 10 to 100 times in 10 mL of selection medium containing 1 μg/mL of puromycin. The cells were seeded in a 10 cm dish, and the selection medium was exchanged every 3 to 4 days. After 8 to 10 days, colonies were marked using a marker pen under a microscope, picked by pipetting with 10 μL of trypsin-EDTA, and subsequently transferred to a 96-well plate containing 100 μL of the selection medium. The cells were allowed to grow until confluency and subcultured 24-well plates and 6-well plates. The cells containing correct PEX5C11A knock-in were identified through PCR genotyping.

### Genomic DNA isolation and PCR

To extract genomic DNA, cells were first lysed in buffer A solution (100 mM Tris-HCl [pH 7.5], 100 mM EDTA, 100 mM NaCl, 0.5% SDS) followed by incubation at 65 °C for 30 min. Buffer B (1.43 M potassium acetate, 4.28 M lithium chloride) was then added and incubated on ice for 10 min. After centrifuging at 12,000 rpm for 15 min, the supernatant was transferred to a new microtube with isopropanol. Precipitated genomic DNA was washed in 70% ethanol and resuspended with DNase-free water. To verify Tet-PEX5C11A-PURO insertion into the AAVS1 locus, genomic PCR was performed using Q5 High-Fidelity DNA polymerase (New England BioLabs). Primers for WT cell validation are: 5′-cgtttcttaggatggccttc-3′ and 5′-agaaggatggagaaagagaa-3′. Primers for Tet-PEX5C11A-PURO integration are: 5′-cgtttcttaggatggccttc-3′ and 5′-ccgggtaaatctccagagga-3′

### Genome wide CRISPR screening

Tet-PEX5^C11A^ HEK293 cells stably expressing human codon-optimized *S.pyogenes* Cas9 were transduced at a multiplicity of infection (MOI) of approximately 0.3 with lentivirus packaged from the Brunello human genome-wide sgRNA library, which contains 76,441 sgRNAs targeting 19,114 protein-coding genes (∼4 sgRNAs per gene), at ≥500-fold sgRNA library representation (*90*). Forty-eight hours of post transduction, cells were selected with puromycin (1 μg/mL) for 7 days to enrich for transduced cells. Following selection, cells were then split into matched Dox−arm (no peroxisomal stress) and Dox+ arm (1 μg/mL doxycycline, peroxisomal stress induced via PEX5C11A expression). Cells underwent three rounds of doxycycline treatment and were propagated for 14-21 days total, while maintaining sgRNA representation above 500× throughout. At harvest, genomic DNA was isolated using the Zymo Quick-DNA Midiprep Plus Kit, sgRNA cassettes were PCR-amplified using barcoded primers, and amplicons were sequenced on an Illumina NextSeq instrument to a depth of reads per sample. Reads were demultiplexed and aligned to the Brunello library reference, and sgRNA counts were analysed using the MAGeCK pipeline(*140*) with default parameters to identify sgRNAs enriched and depleted in Dox+ relative to Dox− conditions. Genes with a fold change > 2 and p < 0.05 by the MAGeCK RRA algorithm were considered significantly enriched or depleted. Screens were performed in independent biological replicates. In total, the screen identified 1,589 sgRNAs corresponding to 1,528 human genes that were significantly enriched or depleted upon peroxisomal stress induction.

### Generation of SCAF1 knockout stable cell line

A stable SCAF1 knockout cell line was generated in HAP1 human haploid cells using CRISPR/Cas9. A chemically modified Alt-R™ sgRNA (IDT; 2 nmol) targeting exon 6 of *SCAF1*; protospacer: 5’-CTGGTGGCTGAGGTCCGAAT-3’) was co-transfected with the pSpCas9(BB)-2A-GFP expression plasmid (PX458; Addgene #48138) in HAP1 cells using Effectene Transfection Reagent (Qiagen; cat. no. 301425) according to the manufacturer’s protocol. At 48 h post-transfection, GFP-positive cells were single-cell sorted into 96-well plates at the Iowa State University Flow Cytometry Facility using a 488 nm laser with a 530/30 nm bandpass filter, gated against untransfected controls. Individual clones were expanded over 10–14 days in conditioned IMDM supplemented with 10% FBS.

Genomic DNA was extracted from expanded clones as described above. The region spanning the exon 6 cut site was amplified by PCR and submitted for Sanger sequencing at the Iowa State University DNA Sequencing Facility. Chromatograms were analyzed using the Snapgene tool to identify indel sequences. Given the near-haploid nature of HAP1 cells, a frameshift indel confirmed by snapgene analysis was considered sufficient to produce a functional null mutation. Induction of MT-ATP6 protein was confirmed as the primary validation criterion for SCAF1 loss by Western blot in at least two independently derived clones, with ATP5A1 as a loading control. Validated knockout clones were used for all downstream experiments. See Key resources table for antibody information

### Cell culture and maintenance

HAP1 human haploid cells (Horizon Discovery) were used for knockout generation. HAP1 cells are derived from KBM7 chronic myelogenous leukaemia cells and are near-haploid for most chromosomes, ensuring that a single allelic disruption produces a complete genetic null without the need for biallelic editing. Cells were maintained in Iscove’s Modified Dulbecco’s Medium (IMDM; Gibco) supplemented with 10% (v/v) fetal bovine serum (FBS; Gibco) and 1% (v/v) penicillin/streptomycin at 37°C in a humidified atmosphere of 5% CO₂.

### Western blotting

Tet-PEX5^C11A^ cells were seeded in 6-well plates. After one day, cells were treated with or without Dox for 3 days. The proteins were extracted in NP-40 cell lysis buffer (Thermo Fisher Scientific, FNN0021) containing 1×protease inhibitor cocktail (Sigma). Protein samples were denatured with Laemmli sample buffer (Bio-Rad, #161-0737) at 95 °C for 5 min. Then proteins were separated by Mini-PROTEAN TGX Precast Gels (Bio-Rad). Following incubation with primary and secondary antibodies, the blots were visualized with Pierce ECL Western Blotting Substrate (Thermo Scientific). See Key resources table for antibody information.

### Cell fitness assay

2.5 × 10^4^ Tet-PEX5^C11A^ cells were seeded in a 96-well plate. Next day, cells were transfected with 20 nM of siRNAs using Opti-MEM (Thermo, 31985062) and RNAiMAX (Thermo, 13778150) according to the manufacturer’s instructions. After 3 hours, cells were incubated with or without Dox (1 μg/mL) for 3 days, and the number of cells was counted manually. All siRNA molecules were obtained from (IDT, Coralville, IA, USA), see Key resources table.

### Mitochondria Complex I Activity Assay

Complex I (NADH:ubiquinone oxidoreductase) activity was measured spectrophotometrically in crude mitochondrial fractions as described by Spinazzi et al. 2012 (*141*), with adaptations for a 96-well plate format. HEK293 Tet-PEX5^C11A^ cells were seeded at 2 × 10⁶ cells per 6-well plate and transfected with scramble control siRNA (Negative Control DsiRNA; IDT, cat. no. 155559763; scramble siRNA) and SCAF1-targeting siRNA (hs.Ri.SCAF1.13.3; IDT, cat. no. 160677271; *SCAF1* siRNA) using Lipofectamine RNAiMAX (Thermo Fisher Scientific) according to the manufacturer’s protocol. Twenty-four hours post-transfection, one plate per condition was treated with doxycycline (1 μg/mL) for 48 h to induce peroxisomal stress via PEX5^C11A^ expression, while the matched control no doxycycline treatment. Cells were washed once with 1 mL ice-cold PBS, scraped and collected by centrifugation at 1,000 × *g* for 5 min at 4°C.

Crude mitochondria were isolated by hypotonic lysis as described by Spinazzi et al. 2012 (*141*). Following a second PBS wash and centrifugation at 1,000 × *g* for 5 min at 4°C, the cell pellet was flash-frozen in liquid nitrogen. For lysis, the frozen pellet was thawed on ice and resuspended in 1ml ice-cold hypotonic 10 mM Tris-HCl (pH 7.6). Cells were disrupted by homogenization in a glass Dounce homogenizer. Isotonicity was restored by addition of 200 μL of 1.5 M sucrose, followed by thorough mixing. The homogenate was centrifuged at 600 × *g* for 10 min at 4°C to remove cell debris and unbroken cells. The supernatant was collected and centrifuged at 14,000 × *g* for 10 min at 4°C to pellet crude mitochondria. The mitochondrial pellet was washed by resuspension in 0.5 mL ice-cold 10 mM Tris-HCl (pH 7.6) and subjected to three freeze-thaw cycles in hypotonic buffer to maximize enzymatic activity, as recommended by Spinazzi et al. [2012](*141*). Protein concentration was determined by BCA assay.

Complex I activity was determined by monitoring rotenone-sensitive NADH oxidation at 340 nm at 30°C in a 96-well plate format. Each reaction contained 50 mM potassium phosphate buffer (pH 7.5), 3.75 mg/mL fatty acid-free BSA, 1 mM KCN (to inhibit Complex IV), and 200 μM NADH, with 20 μg mitochondrial protein per well in a total volume of 200 μL. Reactions were initiated by addition of 100 μM ubiquinone-1 (CoQ₁) and absorbance was read every 1 min for 30 min. Parallel rotenone-containing reactions (rotenone added at 5 μM prior to CoQ₁) were run simultaneously for each sample to determine non-specific NADH oxidation. Rotenone-sensitive Complex I activity was calculated as the difference in the rate of NADH oxidation between the two conditions, expressed as nmol NADH oxidised per minute per mg protein using an extinction coefficient of 6.22 mM⁻¹ cm⁻¹. Three independent biological replicates were performed.

### Mitochondria Isolation

Mitochondria were isolated from cultured cells by differential centrifugation as follows. Cells were grown to 80 % confluency in T-25 flasks, washed once with ice-cold PBS, scraped into 1 mL ice-cold PBS centrifuge at 500g for 3 min. To the pellet 1 ml of hypotonic lysis buffer (10 mM HEPES-KOH pH 7.4, 10 mM KCl, 0.5 mM MgCl₂) was added and incubated on ice for 5 min to promote cell swelling. Cells were lysed by 30 strokes of a glass Dounce homogenizer (Kimble/Kontes, Vineland, NJ, USA), and lysis was confirmed by microscopic inspection of a 5 μL aliquot. To restore isotonicity, 250 μL of sucrose buffer (1 M sucrose, 10 mM HEPES-KOH pH 7.4) was added to achieve a final sucrose concentration of 0.25 M. Cell debris and nuclei were removed by two sequential centrifugations at 1,500×g for 3 min at 4°C, with the supernatant collected after each spin. The post-nuclear supernatant was centrifuged at 10,000×g for 10 min at 4°C; the resulting supernatant was retained as the cytosolic fraction, and the crude mitochondrial pellet was gently washed once with ice-cold PBS. For protein extraction, the crude mitochondria pellet was resuspended in CHAPS lysis buffer (25 mM Tris-HCl pH 8.0, 150 mM NaCl, 2% CHAPS, 1× protease inhibitor cocktail) proportional to the initial pellet weight and incubated for 30 mins. Lysates were clarified by centrifugation at 12,000×g for 2 min, and the supernatant was combined with an equal volume of denaturing sample buffer and heated at 70°C for 10 min and later used for western blot to check protein-level of all the mitochondria encoded proteins.

### Mitochondrial and Cytosolic Fractionation

Mitochondrial and cytosolic fractions were isolated from HEK293 cells using the Mitochondria Isolation Kit for Cultured Cells (Thermo Fisher Scientific, 89874) according to the manufacturer’s protocol. Cells were grown in T-25 flasks and treated with doxycycline (1 μg/mL) for 7 days to induce peroxisomal stress prior to fractionation. Two biological replicates were processed independently. Briefly, cells were washed with 1 mL PBS, scraped into 1 mL fresh PBS, and collected by centrifugation at 500 × *g* for 3 min. The cell pellet was then lysed by reagent-based disruption according to the kit protocol. The lysate was centrifuged at 700 × *g* for 10 min at 4°C to remove cell debris and unbroken cells. The resulting supernatant was further centrifuged at 3,000 × *g* for 15 min at 4°C to pellet the crude mitochondrial fraction, and the supernatant was retained as the cytosolic fraction. The crude mitochondrial pellet was resuspended in CHAPS lysis buffer (25 mM Tris-HCl pH 8.0, 150 mM NaCl, 2% CHAPS, 1× protease inhibitor cocktail) proportional to pellet weight and incubated for 30 min on ice. Lysates were clarified by centrifugation at 12,000 × *g* for 2 min at 4°C, and the supernatant was combined with an equal volume of denaturing sample buffer and heated at 70°C for 10 min prior to Western blot analysis. Protein concentrations of both fractions were determined by BCA assay (Thermo Fisher Scientific). Equal protein amounts were resolved by SDS-PAGE and transferred to a PVDF membrane. Fractionation purity was confirmed to be using ATP5A as a mitochondrial marker and GAPDH as a cytosolic marker. SCAF1 distribution across fractions was subsequently assessed by Western blot; see Key Resources Table for antibody details.

### Immunofluorescence microscopy

HEK293 Tet-PEX5^C11A^ cells were seeded in 24-well plates on poly-L-lysine coated coverslips (Neuvitro #GG1215PLL). Next day, the cells were treated with or without doxycycline (1 μg/mL) for 2 days. The cells were fixed in 4% paraformaldehyde for 10 min, and rinsed with 1× PBS, then permeabilized in 0.5% Triton X-100 in PBS for 10 min. Cells were blocked in PBS containing 1% bovine serum albumin for 1 h at RT, then incubated with anti-SCAF1 (Thermo Fisher Scientific, #10594-1-AP) and anti-ATP5A1 diluted in PBS for overnight at 4 °C. Next day, cells were incubated with secondary antibodies (Alexa Flour 594 donkey anti-rabbit IgG [1:1,000] for 1 h at RT). After washing, cells were mounted using ProLong Gold antifade reagent (Thermo Fisher Scientific) and imaged with an FV3000 Confocal Laser Scanning Microscope (Olympus). Hoechst 33342 was used for nuclear staining.

### Mitochondria Sucrose density Gradient sedimentation

Mitochondria were isolated from Tet-PEX5C^11A^ HEK293, and HAP1 cells grown in T-75 flasks (two flasks per experiment). Next day, the cells were treated with or without doxycycline (1 μg/mL) for 7 days until the cell reaches 80–90% confluency (*95*, *142*). Cells were washed with 4 mL ice-cold PBS, which was discarded. Cells were then scraped into 4 mL fresh ice-cold PBS, transferred to a 15 mL tube, and centrifuged at 500 × *g* for 3 min at 4°C. The pellet was resuspended in 2 mL Mitochondria Isolation Buffer (MIB) (320 mM sucrose, 10 mM Tris-HCl pH 7.5, 1 mM PMSF, 1× protease inhibitor cocktail) and incubated on ice for 5 min. Cells were homogenized in a pre-chilled Dounce homogenizer with 30 strokes per mL (performed in two sequential 1 mL aliquots). The homogenate was centrifuged at 1,000 × *g* for 10 min at 4°C to remove cell debris and unbroken cells. The supernatant was transferred to a new tube and centrifuged at 12,000 × *g* for 10 min at 4°C to pellet crude mitochondria. The mitochondrial pellet was washed once by resuspension in 1 mL ice-cold MIB and re-centrifuged at 12,000 × *g* for 10 min at 4°C. Protein concentration was determined by BCA assay prior to lysis For sucrose gradient sedimentation, the mitochondrial pellet was resuspended in mitochondria lysis buffer (MLB: 260 mM sucrose, 100 mM KCl, 20 mM MgCl₂, 10 mM Tris-HCl pH 7.5, 1% Triton X-100, 5 mM β-mercaptoethanol, 1 mM PMSF, 1× protease inhibitor cocktail) at a volume proportional to protein concentration as determined by BCA, targeting equal protein input across samples. Lysates were incubated on ice for 20 min with periodic mixing, then clarified by centrifugation at 9,200 × *g* for 45 min at 4°C. 10–30% linear sucrose gradients were prepared in gradient buffer (100 mM KCl, 20 mM MgCl₂, 10 mM Tris-HCl pH 7.5, 5 mM β-mercaptoethanol, 1 mM PMSF, 1× protease inhibitor cocktail) using a Gradient Master (BioComp Instruments, Inc.) in SW 55 Ti ultracentrifuge tubes. Prior to sample loading, 200 μL was removed from the top of each gradient. Clarified mitochondrial lysate (350 μL) was gently layered onto each gradient and centrifuged at 40,000 × *g* for 3 h 30 min at 4°C in an SW 55 Ti rotor (Beckman Coulter) with maximum acceleration and minimum deceleration. Twenty-four fractions of 180 μL each were collected from the top using a BioComp Piston Gradient Fractionator. Proteins from each fraction were resolved by SDS-PAGE and analyzed by Western blot using antibodies against SCAF1, MRPL12, MRPL11, MRPS15, WBSCR16, LRPPRC, FASTKD4, and GTPBP10 (1:2000 dilution).

### Co-immunoprecipitation

HAP1 cells were transfected with truncated S1 - pCMV-MTS Cox8-SCAF1-1-205-FLAG; S2 - pCMV-MTS Cox8-SCAF1-205-425-FLAG; S5-pCMV-MTS Cox8-SCAF1-887-1087-FLAG; S6-pCMV-MTS Cox8 SCAF1-1088-1312-FLAG in HAP1 cells using Lipofectamine RNAiMAx after 48 hrs. Whole-cell lysates were prepared from HAP1 cells in IP lysis buffer (50 mM Tris-HCl pH 7.5, 150 mM NaCl, 1% NP-40, 0.5% sodium deoxycholate, 1× protease inhibitor cocktail). Lysates were precleared with Dyna beads for 1 h at 4 °C and then incubated with anti-Flag antibody (Sigma #F1804) 10 μg overnight at 4 °C. Antibody-antigen complexes were captured with Dynabeads Protein G for 2 h at 4 °C, washed the beads three times in IP lysis buffer, eluted with Laemmli buffer at 95 °C for 5 min. And immunoblot with WBSCR16, GTPBP10, DDX28, DHX30, FASTKD4, FASTKD2, ERAL1, MALSU1 and METL17A.

### Senescence-Associated β-Galactosidase Staining

To evaluate cellular senescence following *PEX5* knockdown, and MTS Knocdown senescence-associated β-galactosidase (SA-β-gal) activity was determined using the Senescence β-Galactosidase Staining Kit #9860 from Cell Signaling Technology. Following the experimental incubation, IMR90 cells were rinsed once with 1× phosphate-buffered saline (PBS) and treated with the provided 1× fixative solution for 10 to 15 min at room temperature. Fixed cells were washed twice with 1× PBS and then incubated overnight (16 to 24 hours) at 37 °C in a dry incubator without CO₂ with a freshly prepared, pH 6.0 development solution containing 5-bromo-4-chloro-3-indolyl-β-D-galactopyranoside (X-gal). The enzymatic staining reaction was terminated by rinsing the monolayers twice with 1× PBS. Blue-stained senescent cells were visualized, and representative images were captured using a bright-field microscope. The proportion of SA-β-gal–positive cells were quantified by scoring at least five random fields per experimental group to evaluate the total percentage of senescent cells.

### Quantitative RT-PCR

Total RNA was extracted using TRIzol (Thermo Fisher Scientific) and reverse-transcribed using the High-Capacity cDNA Reverse Transcription Kit (Applied Biosystems). Quantitative PCR was performed using SYBR Green PCR Master Mix (Applied Biosystems) on a QuantStudio 6 instrument. Mitochondrial transcripts were normalised to nuclear-encoded ACTB. Primer sequences are listed in the Key Resources Table. P-values for multi-gene comparisons (Figure 3A) were corrected for multiple testing using the Benjamini–Hochberg false discovery rate (q < 0.05).

### Statistical Analysis

GraphPad Prism (GraphPad Software, La Jolla, CA, USA) was used for statistical analysis. Detailed tests used are given in the corresponding figure legends. Statistical analysis was performed using either an unpaired two-tailed t test or one-way ANOVA with Tukey multiple comparison.

**Table 1:**
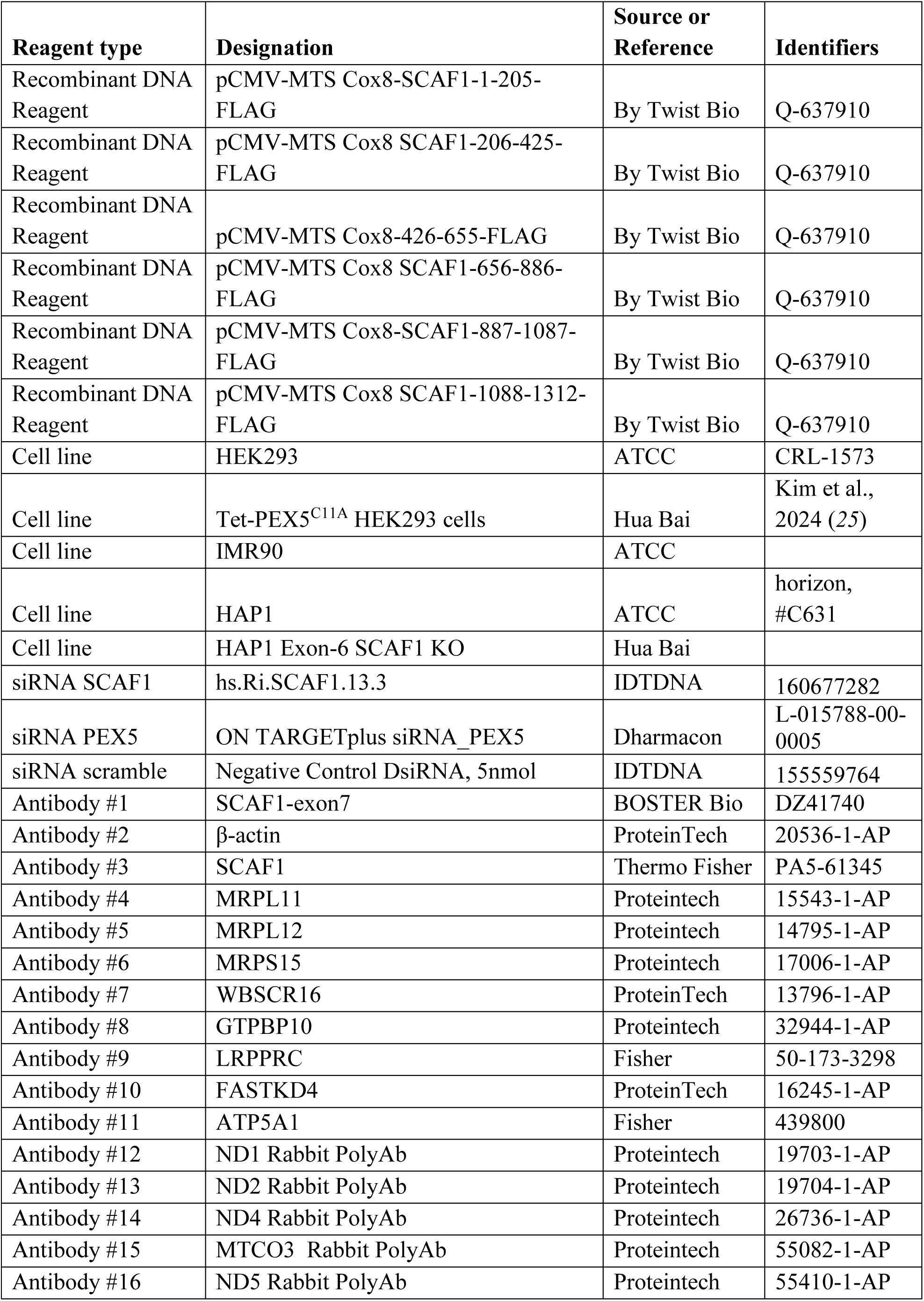
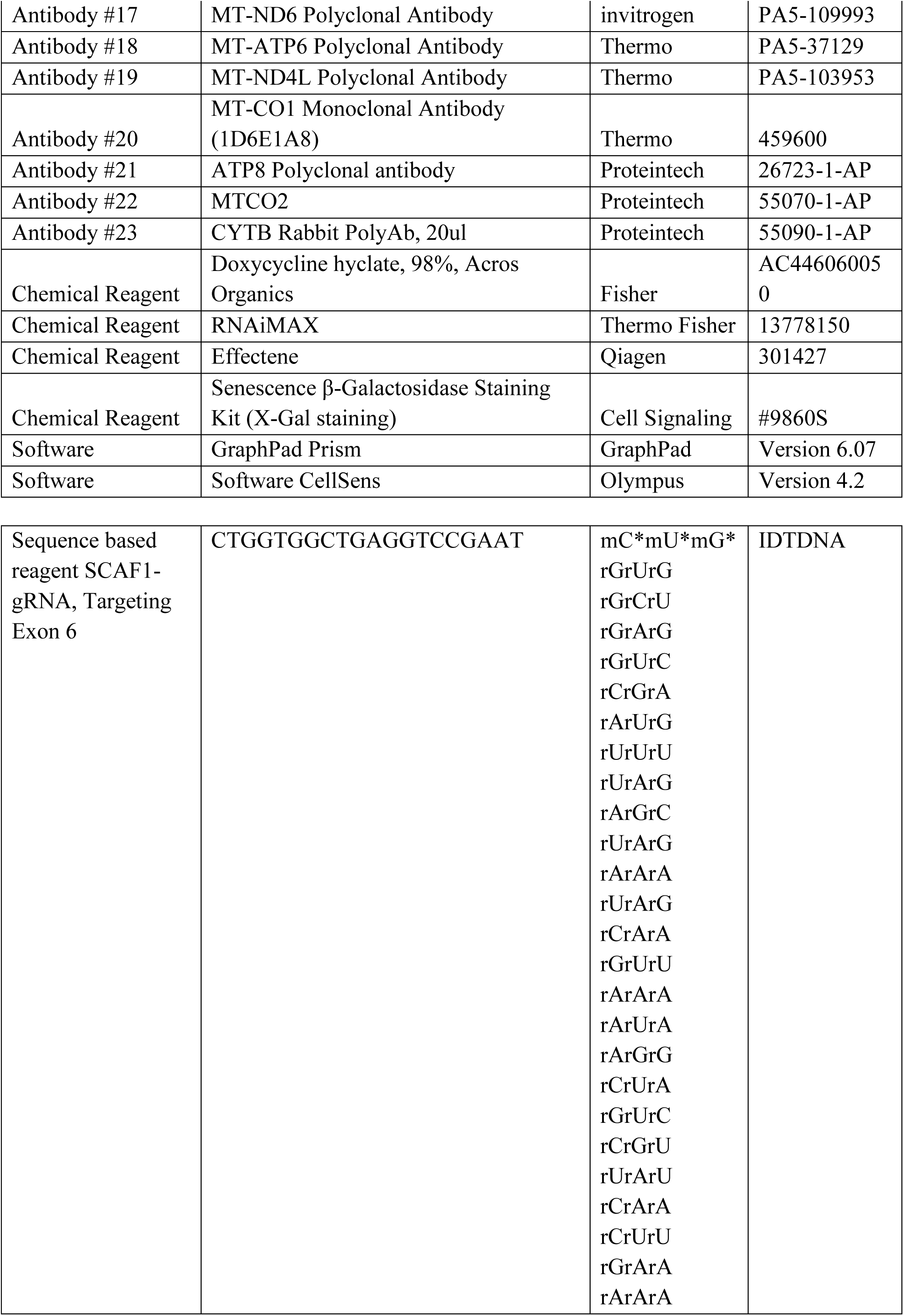
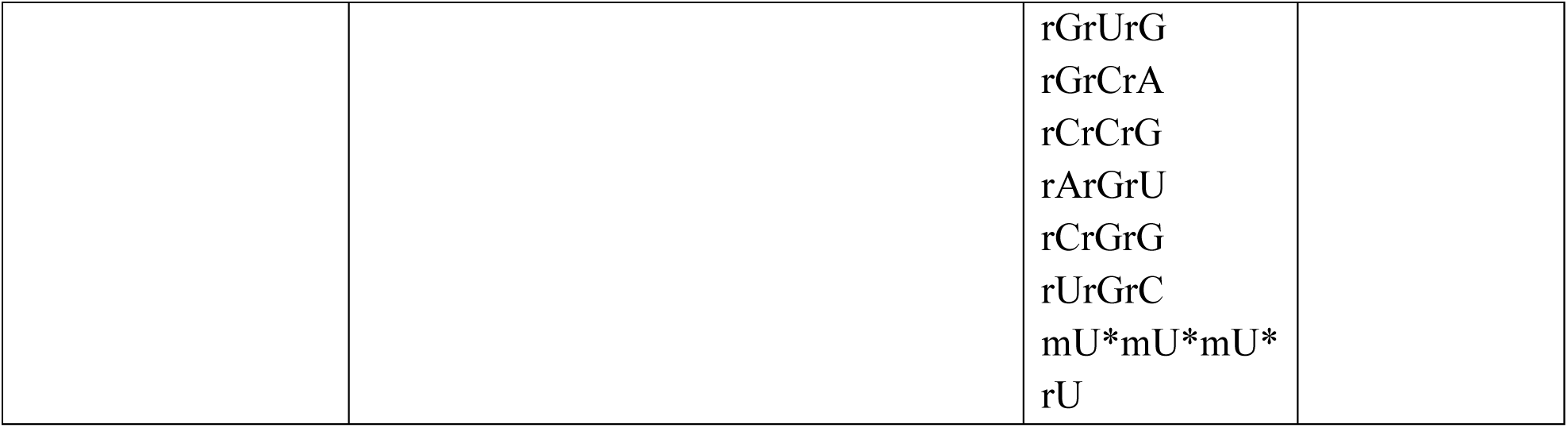
List of Plasmid, Reagents and Antibody.

**Table 2:**
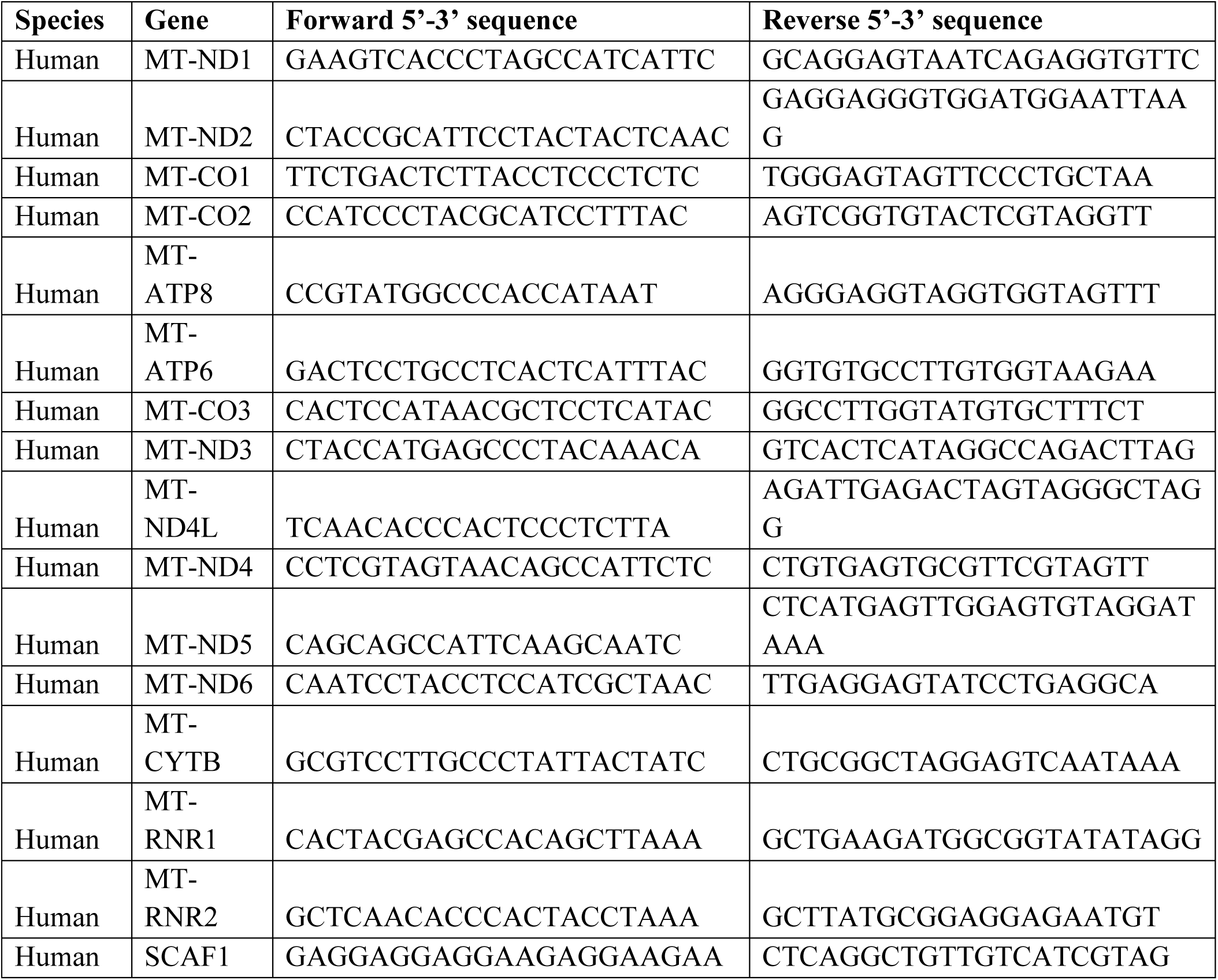
List of Primers.

## Acknowledgments

We thank Iowa State University for institutional support. This work was supported by National Science Foundation (NSF) CAREER 2046984 awarded to Hua Bai. We thank the ISU DNA sequencing Facility.

## Funding

National Institute of Health (NIH) R01AG058741 to H.B.

## Author contributions

Conceptualization: J.C., J.K., H.B.

Methodology: J.C., J.K., H.B.

Investigation: J.C., J.K., P.V.

Visualization: J.C., J.K.

Supervision: J.C., J.K., H.B.

Writing—original draft: J.C., H.B

Writing—review & editing: J.C., H.B.

## Competing interests

The authors have no competing interest

---

**Supplementary Fig. 1.**
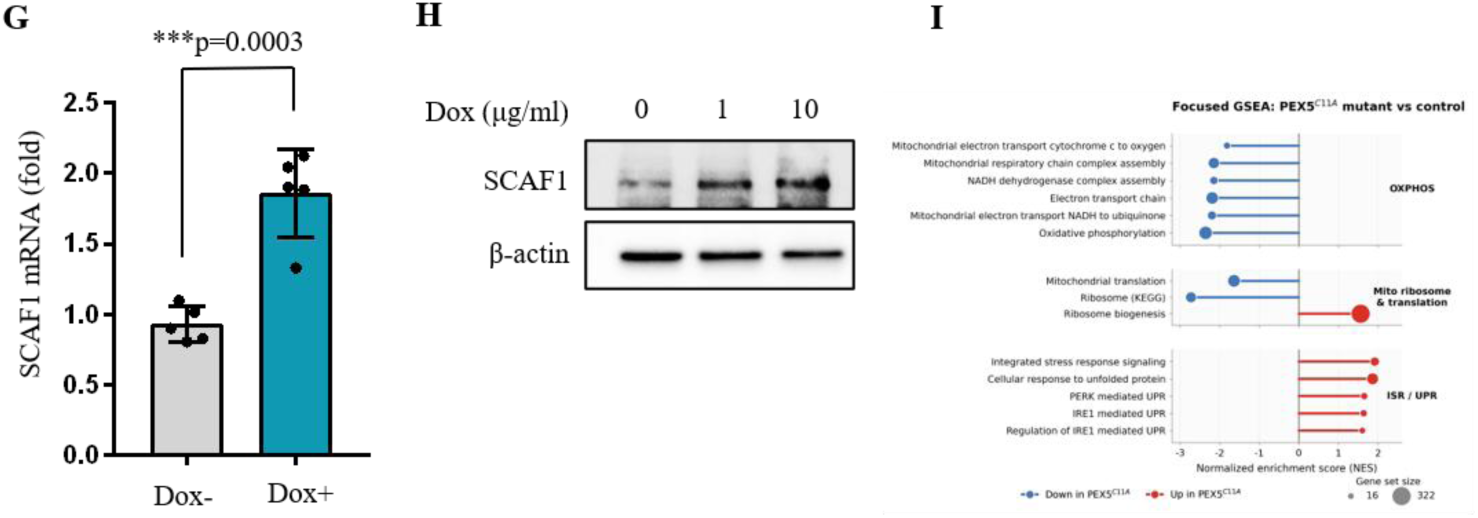
CRISPR screen identify SCAF1 as a regulator mediating peroxisome stress. (G) SCAF1 mRNA expression was measured by qPCR after 3 days of doxycycline treatment (1μg/ml) on Tet-On PEX5^C11A^ cells. ***p =0.0003, mean ± SD, t-test. (H) SCAF1 protein expression was measured by western blotting after 3 days of doxycycline treatment (0, 1, or 10 μg/ml) on Tet-ON PEX5^C11A^ cells. (I). Gene set Enrichment Analysis of Mitochondrial and stress response pathways in Tet-On PEX5^C11A^ cells. Bar chart showing normalized enrichment scores (NES) for significantly enriched gene sets ( padj<0.05) across KEGG, HALLMARK and GO Biological Process (GO:BP) databases. Bars extending left (blue) indicate downregulation in PEX5 ^C11A^ mutant relative to control; bars extending right (red) indicate upregulation. GSEA was performed using the fgsea package in R with gene sets from MSigDB.

## Notes

### Competing Interest Statement

The authors have declared no competing interest.

## REFERENCES

1. López-Otín, C., et al., Hallmarks of aging: An expanding universe. Cell, 2023. 186(2): p. 243–278.

2. Moldakozhayev, A. and V.N. Gladyshev, Metabolism, homeostasis, and aging. Trends Endocrinol Metab, 2023. 34(3): p. 158–169.

3. Amorim, J.A., et al., Mitochondrial and metabolic dysfunction in ageing and age-related diseases. Nat Rev Endocrinol, 2022. 18(4): p. 243–258.

4. Finkel, T., The metabolic regulation of aging. Nat Med, 2015. 21(12): p. 1416–23.

5. Mironova, E., et al., Mitochondria and Aging: Redox Balance Modulation as a New Approach to the Development of Innovative Geroprotectors (Fundamental and Applied Aspects). Int J Mol Sci, 2026. 27(2).

6. Miwa, S., et al., Mitochondrial dysfunction in cell senescence and aging. J Clin Invest, 2022. 132(13).

7. Hruby, A.J. and R. Higuchi-Sanabria, Mitochondrial dysfunction in cellular senescence: a bridge to neurodegenerative disease. NPJ Aging, 2025. 11(1): p. 99.

8. Martini, H. and J.F. Passos, Cellular senescence: all roads lead to mitochondria. Febs j, 2023. 290(5): p. 1186–1202.

9. Chen, B., C.A. Lyssiotis, and Y.M. Shah, Mitochondria-organelle crosstalk in establishing compartmentalized metabolic homeostasis. Mol Cell, 2025. 85(8): p. 1487–1508.

10. J. Kim, H. Bai, Peroxisomal Stress Response and Inter-Organelle Communication in Cellular Homeostasis and Aging. Antioxidants 11, 192 (2022).

11. J. J. Smith, J. D. Aitchison, Peroxisomes take shape. Nat Rev Mol Cell Biol 14, 803–817 (2013).

12. T. Walter, R. Erdmann, Current Advances in Protein Import into Peroxisomes. Protein J 38, 351–362 (2019).

13. M. Fransen, C. Lismont, P. Walton, The Peroxisome-Mitochondria Connection: How and Why? IJMS 18, 1126 (2017).

14. R. J. A. Wanders, H. R. Waterham, Biochemistry of Mammalian Peroxisomes Revisited. Annu. Rev. Biochem. 75, 295–332 (2006).

15. H. R. Waterham, S. Ferdinandusse, R. J. A. Wanders, Human disorders of peroxisome metabolism and biogenesis. Biochimica et Biophysica Acta (BBA) - Molecular Cell Research 1863, 922–933 (2016).

16. M. Schrader, H. D. Fahimi, The peroxisome: still a mysterious organelle. Histochem Cell Biol 129, 421–440 (2008).

17. S. J. Steinberg, G. Dodt, G. V. Raymond, N. E. Braverman, A. B. Moser, H. W. Moser, Peroxisome biogenesis disorders. Biochimica et Biophysica Acta (BBA) - Molecular Cell Research 1763, 1733–1748 (2006).

18. F. C. C. Klouwer, K. Berendse, S. Ferdinandusse, R. J. A. Wanders, M. Engelen, B. T. Poll-The, Zellweger spectrum disorders: clinical overview and management approach. Orphanet J Rare Dis 10, 151 (2015).

19. A. F. Carvalho, M. P. Pinto, C. P. Grou, I. S. Alencastre, M. Fransen, C. Sá-Miranda, J. E. Azevedo, Ubiquitination of Mammalian Pex5p, the Peroxisomal Import Receptor. Journal of Biological Chemistry 282, 31267–31272 (2007).

20. C. Williams, M. Van Den Berg, R. R. Sprenger, B. Distel, A Conserved Cysteine Is Essential for Pex4p-dependent Ubiquitination of the Peroxisomal Import Receptor Pex5p. Journal of Biological Chemistry 282, 22534–22543 (2007).

21. H. W. Platta, F. El Magraoui, B. E. Bäumer, D. Schlee, W. Girzalsky, R. Erdmann, Pex2 and Pex12 Function as Protein-Ubiquitin Ligases in Peroxisomal Protein Import. Molecular and Cellular Biology 29, 5505–5516 (2009).

22. K. Okumoto, S. Misono, N. Miyata, Y. Matsumoto, S. Mukai, Y. Fujiki, Cysteine Ubiquitination of PTS1 Receptor Pex5p Regulates Pex5p Recycling. Traffic 12, 1067–1083 (2011).

23. K. Huang, T. Miao, K. Chang, J. Kim, P. Kang, Q. Jiang, A. J. Simmonds, F. Di Cara, H. Bai, Impaired peroxisomal import in Drosophila oenocytes causes cardiac dysfunction by inducing upd3 as a peroxikine. Nat Commun 11, 2943 (2020).

24. J. Correia, P. S. Roy, K. G. Holden, M. L. Kohut, H. Bai, Aging impairs peroxisome biogenesis in human B cells. *The Journals of Gerontology*, Series A: Biological Sciences and Medical Sciences 81, glaf148 (2026).

25. J. Kim, K. Huang, P. T. T. Vo, T. Miao, J. Correia, A. Kumar, M. J. P. Simons, H. Bai, Peroxisomal import stress activates integrated stress response and inhibits ribosome biogenesis. PNAS Nexus 3, pgae429 (2024).

26. E. Baumgart, I. Vanhorebeek, M. Grabenbauer, M. Borgers, P. E. Declercq, H. D. Fahimi, M. Baes, Mitochondrial Alterations Caused by Defective Peroxisomal Biogenesis in a Mouse Model for Zellweger Syndrome (PEX5 Knockout Mouse). The American Journal of Pathology 159, 1477–1494 (2001).

27. R. S. Rahim, M. Chen, C. C. Nourse, A. C. B. Meedeniya, D. I. Crane, Mitochondrial changes and oxidative stress in a mouse model of Zellweger syndrome neuropathogenesis. Neuroscience 334, 201–213 (2016).

28. M. Schrader, N. A. Bonekamp, M. Islinger, Fission and proliferation of peroxisomes. Biochimica et Biophysica Acta (BBA) - Molecular Basis of Disease 1822, 1343–1357 (2012).

29. N. Shai, E. Yifrach, C. W. T. Van Roermund, N. Cohen, C. Bibi, L. IJlst, L. Cavellini, J. Meurisse, R. Schuster, L. Zada, M. C. Mari, F. M. Reggiori, A. L. Hughes, M. Escobar-Henriques, M. M. Cohen, H. R. Waterham, R. J. A. Wanders, M. Schuldiner, E. Zalckvar, Systematic mapping of contact sites reveals tethers and a function for the peroxisome-mitochondria contact. Nat Commun 9, 1761 (2018).

30. J. L. Costello, I. G. Castro, C. Hacker, T. A. Schrader, J. Metz, D. Zeuschner, A. S. Azadi, L. F. Godinho, V. Costina, P. Findeisen, A. Manner, M. Islinger, M. Schrader, ACBD5 and VAPB mediate membrane associations between peroxisomes and the ER. Journal of Cell Biology 216, 331–342 (2017).

31. R. Hua, D. Cheng, É. Coyaud, S. Freeman, E. Di Pietro, Y. Wang, A. Vissa, C. M. Yip, G. D. Fairn, N. Braverman, J. H. Brumell, W. S. Trimble, B. Raught, P. K. Kim, VAPs and ACBD5 tether peroxisomes to the ER for peroxisome maintenance and lipid homeostasis. Journal of Cell Biology 216, 367–377 (2017).

32. B. J. Greber, D. Boehringer, M. Leibundgut, P. Bieri, A. Leitner, N. Schmitz, R. Aebersold, N. Ban, The complete structure of the large subunit of the mammalian mitochondrial ribosome. Nature 515, 283–286 (2014).

33. A. Amunts, A. Brown, J. Toots, S. H. W. Scheres, V. Ramakrishnan, The structure of the human mitochondrial ribosome. Science 348, 95–98 (2015).

34. A. Brown, A. Amunts, X. Bai, Y. Sugimoto, P. C. Edwards, G. Murshudov, S. H. W. Scheres, V. Ramakrishnan, Structure of the large ribosomal subunit from human mitochondria.

35. A. Brown, S. Rathore, D. Kimanius, S. Aibara, X. Bai, J. Rorbach, A. Amunts, V. Ramakrishnan, Structures of the human mitochondrial ribosome in native states of assembly. Nat Struct Mol Biol 24, 866–869 (2017).

36. D. De Silva, Y.-T. Tu, A. Amunts, F. Fontanesi, A. Barrientos, Mitochondrial ribosome assembly in health and disease. Cell Cycle 14, 2226–2250 (2015).

37. S. F. Pearce, P. Rebelo-Guiomar, A. R. D’Souza, C. A. Powell, L. Van Haute, M. Minczuk, Regulation of Mammalian Mitochondrial Gene Expression: Recent Advances. Trends in Biochemical Sciences 42, 625–639 (2017).

38. D. F. Bogenhagen, D. W. Martin, A. Koller, Initial Steps in RNA Processing and Ribosome Assembly Occur at Mitochondrial DNA Nucleoids. Cell Metabolism 19, 618–629 (2014).

39. D. F. Bogenhagen, A. G. Ostermeyer-Fay, J. D. Haley, M. Garcia-Diaz, Kinetics and Mechanism of Mammalian Mitochondrial Ribosome Assembly. Cell Reports 22, 1935–1944 (2018).

40. J. Rorbach, M. Minczuk, The post-transcriptional life of mammalian mitochondrial RNA. Biochemical Journal 444, 357–373 (2012).

41. E. Kummer, N. Ban, Mechanisms and regulation of protein synthesis in mitochondria. Nat Rev Mol Cell Biol 22, 307–325 (2021).

42. M. Cipullo, G. V. Gesé, A. Khawaja, B. M. Hällberg, J. Rorbach, Structural basis for late maturation steps of the human mitoribosomal large subunit. Nat Commun 12, 3673 (2021).

43. P. Maiti, H.-J. Kim, Y.-T. Tu, A. Barrientos, Human GTPBP10 is required for mitoribosome maturation. Nucleic Acids Research, doi: 10.1093/nar/gky938 (2018).

44. E. Lavdovskaia, E. Kolander, E. Steube, M. M.-Q. Mai, H. Urlaub, R. Richter-Dennerlein, The human Obg protein GTPBP10 is involved in mitoribosomal biogenesis. Nucleic Acids Research 46, 8471–8482 (2018).

45. T. Kotani, S. Akabane, K. Takeyasu, T. Ueda, N. Takeuchi, Human G-proteins, ObgH1 and Mtg1, associate with the large mitochondrial ribosome subunit and are involved in translation and assembly of respiratory complexes. Nucleic Acids Research 41, 3713–3722 (2013).

46. H. S. Hillen, E. Lavdovskaia, F. Nadler, E. Hanitsch, A. Linden, K. E. Bohnsack, H. Urlaub, R. Richter-Dennerlein, Structural basis of GTPase-mediated mitochondrial ribosome biogenesis and recycling. Nat Commun 12, 3672 (2021).

47. P. Maiti, E. Lavdovskaia, A. Barrientos, R. Richter-Dennerlein, Role of GTPases in Driving Mitoribosome Assembly. Trends in Cell Biology 31, 284–297 (2021).

48. J. Cheng, O. Berninghausen, R. Beckmann, A distinct assembly pathway of the human 39S late pre-mitoribosome. Nat Commun 12, 4544 (2021).

49. T. G. Nguyen, C. Ritter, E. Kummer, Structural insights into the role of GTPBP10 in the RNA maturation of the mitoribosome. Nat Commun 14, 7991 (2023).

50. H. Antonicka, E. A. Shoubridge, Mitochondrial RNA Granules Are Centers for Posttranscriptional RNA Processing and Ribosome Biogenesis. Cell Reports 10, 920–932 (2015).

51. Y.-T. Tu, A. Barrientos, The Human Mitochondrial DEAD-Box Protein DDX28 Resides in RNA Granules and Functions in Mitoribosome Assembly. Cell Reports 10, 854–864 (2015).

52. F. Sasarman, C. Brunel-Guitton, H. Antonicka, T. Wai, E. A. Shoubridge, LSFC Consortium, LRPPRC and SLIRP Interact in a Ribonucleoprotein Complex That Regulates Posttranscriptional Gene Expression in Mitochondria. MBoC 21, 1315–1323 (2010).

53. B. Ruzzenente, M. D. Metodiev, A. Wredenberg, A. Bratic, C. B. Park, Y. Cámara, D. Milenkovic, V. Zickermann, R. Wibom, K. Hultenby, H. Erdjument-Bromage, P. Tempst, U. Brandt, J. B. Stewart, C. M. Gustafsson, N. Larsson, LRPPRC is necessary for polyadenylation and coordination of translation of mitochondrial mRNAs. EMBO J 31, 443– 456 (2012).

54. S. Fung, T. Nishimura, F. Sasarman, E. A. Shoubridge, The conserved interaction of C7orf30 with MRPL14 promotes biogenesis of the mitochondrial large ribosomal subunit and mitochondrial translation. MBoC 24, 184–193 (2013).

55. J. Rorbach, P. Boesch, P. A. Gammage, T. J. J. Nicholls, S. F. Pearce, D. Patel, A. Hauser, F. Perocchi, M. Minczuk, MRM2 and MRM3 are involved in biogenesis of the large subunit of the mitochondrial ribosome. MBoC 25, 2542–2555 (2014).

56. A. A. Jourdain, M. Koppen, M. Wydro, C. D. Rodley, R. N. Lightowlers, Z. M. Chrzanowska-Lightowlers, J.-C. Martinou, GRSF1 Regulates RNA Processing in Mitochondrial RNA Granules. Cell Metabolism 17, 399–410 (2013).

57. A. A. Jourdain, J. Popow, M. A. de la Fuente, J.-C. Martinou, P. Anderson, M. Simarro, The FASTK family of proteins: emerging regulators of mitochondrial RNA biology. Nucleic Acids Research 45, 10941–10947 (2017).

58. T. R. Richman, H. Spåhr, J. A. Ermer, S. M. K. Davies, H. M. Viola, K. A. Bates, J. Papadimitriou, L. C. Hool, J. Rodger, N.-G. Larsson, O. Rackham, A. Filipovska, Loss of the RNA-binding protein TACO1 causes late-onset mitochondrial dysfunction in mice. Nat Commun 7, 11884 (2016).

59. J. A. Letts, L. A. Sazanov, Clarifying the supercomplex: the higher-order organization of the mitochondrial electron transport chain. Nat Struct Mol Biol 24, 800–808 (2017).

60. I. Vercellino, L. A. Sazanov, The assembly, regulation and function of the mitochondrial respiratory chain. Nat Rev Mol Cell Biol 23, 141–161 (2022).

61. A. M. Zahler, W. S. Lane, J. A. Stolk, M. B. Roth, SR proteins: a conserved family of pre-mRNA splicing factors. Genes Dev. 6, 837–847 (1992).

62. J. C. Long, J. F. Caceres, The SR protein family of splicing factors: master regulators of gene expression. Biochemical Journal 417, 15–27 (2009).

63. J. L. Manley, A. R. Krainer, A rational nomenclature for serine/arginine-rich protein splicing factors (SR proteins): Table 1. Genes Dev. 24, 1073–1074 (2010).

64. P. J. Shepard, K. J. Hertel, The SR protein family. Genome Biol 10, 242 (2009).

65. T. Misteli, J. F. Cáceres, D. L. Spector, The dynamics of a pre-mRNA splicing factor in living cells. Nature 387, 523–527 (1997).

66. S. Stamm, Regulation of Alternative Splicing by Reversible Protein Phosphorylation. Journal of Biological Chemistry 283, 1223–1227 (2008).

67. A. Scorilas, L. Kyriakopoulou, D. Katsaros, E. P. Diamandis, Cloning of a gene (SR-A1), encoding for a new member of the human Ser/Arg-rich family of pre-mRNA splicing factors: overexpression in aggressive ovarian cancer. Br J Cancer 85, 190–198 (2001).

68. P. G. Adamopoulos, G. D. Raptis, C. K. Kontos, A. Scorilas, Discovery and expression analysis of novel transcripts of the human SR-related CTD-associated factor 1 (SCAF1) gene in human cancer cells using Next-Generation Sequencing. Gene 670, 155–165 (2018).

69. M. E. Katsarou, A. Papakyriakou, N. Katsaros, A. Scorilas, Expression of the C-terminal domain of novel human SR-A1 protein: Interaction with the CTD domain of RNA polymerase II. Biochemical and Biophysical Research Communications 334, 61–68 (2005).

70. S. Martinez, S. Wu, M. Geuenich, A. Malik, R. Weber, T. Woo, A. Zhang, G. H. Jang, D. Dervovic, K. N. Al-Zahrani, R. Tsai, N. Fodil, P. Gros, S. Gallinger, G. G. Neely, F. Notta, A. Sendoel, K. Campbell, U. Elling, D. Schramek, In vivo CRISPR screens reveal SCAF1 and USP15 as drivers of pancreatic cancer. Nat Commun 15, 5266 (2024).

71. S. Kompocholi, N. Stamidis, H. Liu, G. M. Hjortø, E. Kafkia, 1 SCAF1 driven polyadenylation site usage regulates 2 mRNA isoform expression and neuronal 3 differentiation.

72. J. Van Setten, J. A. Brody, Y. Jamshidi, B. R. Swenson, A. M. Butler, H. Campbell, F. M. Del Greco, D. S. Evans, Q. Gibson, D. F. Gudbjartsson, K. F. Kerr, B. P. Krijthe, L.-P. Lyytikäinen, C. Müller, M. Müller-Nurasyid, I. M. Nolte, S. Padmanabhan, M. D. Ritchie, A. Robino, A. V. Smith, M. Steri, T. Tanaka, A. Teumer, S. Trompet, S. Ulivi, N. Verweij, X. Yin, D. O. Arnar, F. W. Asselbergs, J. S. Bader, J. Barnard, J. Bis, S. Blankenberg, E. Boerwinkle, Y. Bradford, B. M. Buckley, M. K. Chung, D. Crawford, M. Den Hoed, J. C. Denny, A. F. Dominiczak, G. B. Ehret, M. Eijgelsheim, P. T. Ellinor, S. B. Felix, O. H. Franco, L. Franke, T. B. Harris, H. Holm, G. Ilaria, A. Iorio, M. Kähönen, I. Kolcic, J. A. Kors, E. G. Lakatta, L. J. Launer, H. Lin, H. J. Lin, R. J. F. Loos, S. A. Lubitz, P. W. Macfarlane, J. W. Magnani, I. M. Leach, T. Meitinger, B. D. Mitchell, T. Munzel, G. J. Papanicolaou, A. Peters, A. Pfeufer, P. P. Pramstaller, O. T. Raitakari, J. I. Rotter, I. Rudan, N. J. Samani, D. Schlessinger, C. T. Silva Aldana, M. F. Sinner, J. D. Smith, H. Snieder, E. Z. Soliman, T. D. Spector, D. J. Stott, K. Strauch, K. V. Tarasov, U. Thorsteinsdottir, A. G. Uitterlinden, D. R. Van Wagoner, U. Völker, H. Völzke, M. Waldenberger, H. Jan Westra, P. S. Wild, T. Zeller, A. Alonso, C. L. Avery, S. Bandinelli, E. J. Benjamin, F. Cucca, M. Dörr, L. Ferrucci, P. Gasparini, V. Gudnason, C. Hayward, S. R. Heckbert, A. A. Hicks, J. W. Jukema, S. Kääb, T. Lehtimäki, Y. Liu, P. B. Munroe, A. Parsa, O. Polasek, B. M. Psaty, D. M. Roden, R. B. Schnabel, G. Sinagra, K. Stefansson, B. H. Stricker, P. Van Der Harst, C. M. Van Duijn, J. F. Wilson, S. A. Gharib, P. I. W. De Bakker, A. Isaacs, D. E. Arking, N. Sotoodehnia, PR interval genome-wide association meta-analysis identifies 50 loci associated with atrial and atrioventricular electrical activity. Nat Commun 9, 2904 (2018).

73. X. Zhou, P. Feliciano, C. Shu, T. Wang, I. Astrovskaya, J. B. Hall, J. U. Obiajulu, J. R. Wright, S. C. Murali, S. X. Xu, L. Brueggeman, T. R. Thomas, O. Marchenko, C. Fleisch, S. D. Barns, L. G. Snyder, B. Han, T. S. Chang, T. N. Turner, W. T. Harvey, A. Nishida, B. J. O’Roak, D. H. Geschwind, The SPARK Consortium, A. Adams, A. Amatya, A. Andrus, A. Bashar, A. Berman, A. Brown, A. Camba, A. C. Gulsrud, A. D. Krentz, A. D. Shocklee, A. Esler, A. E. Lash, A. Fanta, A. Fatemi, A. Fish, A. Goler, A. Gonzalez, A. Gutierrez, A. Hardan, A. Hess, A. Hirshman, A. Holbrook, A. J. Ace, A. J. Griswold, A. J. Gruber, A. Jarratt, A. Jelinek, A. Jorgenson, A. P. Juarez, A. Kim, A. Kitaygorodsky, A. Luo, A. L. Rachubinski, A. L. Wainer, A. M. Daniels, A. Mankar, A. Mason, A. Miceli, A. Milliken, A. Morales-Lara, A. N. Stephens, A. N. Nguyen, A. Nicholson, A. M. Paolicelli, A. P. McKenzie, A. R. Gupta, A. Raven, A. Rhea, A. Simon, A. Soucy, A. Swanson, A. Sziklay, A. Tallbull, A. Tesng, A. Ward, A. Zick, B. A. Hilscher, B. Bell, B. Enright, B. E. Robertson, B. Hauf, B. Jensen, B. Lobisi, B. M. Vernoia, B. Schwind, B. VanMetre, C. A. Erickson, C. A. W. Sullivan, C. Albright, C. Anglo, C. Buescher, C. C. Bradley, C. Campo-Soria, C. Cohen, C. Colombi, C. Diggins, C. Edmonson, C. E. Rice, C. Fassler, C. Gray, C. Gunter, C. H. Walston, C. Klaiman, C. Leonczyk, C. L. Martin, C. Lord, C. M. Taylor, C. McCarthy, C. Ochoa-Lubinoff, C. Ortiz, C. Pierre, C. R. Rosenberg, C. Rigby, C. Roche, C. Shrier, C. Smith, C. Van Wade, C. White-Lehman, C. Zaro, C. Zha, D. Bentley, D. Correa, D. E. Sarver, D. Giancarla, D. G. Amaral, D. Howes, D. Istephanous, D. L. Coury, D. Li, D. Limon, D. Limpoco, D. Phillips, D. Rambeck, D. Rojas, D. Srishyla, D. Stamps, D. V. Montes, D. Cho, D. Cho, E. A. Fox, E. Bahl, E. Berry-Kravis, E. Blank, E. Bower, E. Brooks, E. Courchesne, E. Dillon, E. Doyle, E. Given, E. Grimes, E. Jones, E. J. Fombonne, E. Kryszak, E. L. Wodka, E. Lamarche, E. Lampert, E. M. Butter, E. O’Connor, E. Ocampo, E. Orrick, E. Perez, E. Ruzzo, E. Singer, E. T. Matthews, E. V. Pedapati, F. Fazal, F. K. Miller, G. Aberbach, G. Baraghoshi, G. Duhon, G. Hooks, G. J. Fischer, G. Marzano, G. Schoonover, G. S. Dichter, G. Tiede, H. Cottrell, H. E. Kaplan, H. Ghina, H. Hutter, H. Koene, H. L. Schneider, H. Lechniak, H. Li, H. Morotti, H. Qi, H. Richardson, H. Zaydens, H. Zhang, H. Zhao, I. Arriaga, I. F. Tso, J. Acampado, J. A. Gerdts, J. Beeson, J. Brown, J. Comitre, J. Cordova, J. Delaporte, J. F. Cubells, J. F. Harris, J. Gong, J. Gunderson, J. Hernandez, J. Judge, J. Jurayj, J. K. Law, J. Manoharan, J. Montezuma, J. Neely, J. Orobio, J. Pandey, J. Piven, J. Polanco, J. Polite, J. Rosewater, J. Scherr, J. S. Sutcliffe, J. T. McCracken, J. Tjernagel, J. Toroney, J. Veenstra-Vanderweele, J. Wang, K. Ahlers, K. A. Schweers, K. Baalman, K. Beard, K. Callahan, K. Coleman, K. D. Fitzgerald, K. Dent, K. Diehl, K. Gonring, K. G. Pawlowski, K. Hirst, K. L. Pierce, K. Murillo, K. Murray, K. Nowell, K. O’Brien, K. Pama, K. Real, K. Singer, K. Smith, K. Stephenson, K. Tsai, L. Abbeduto, L. A. Cartner, L. Beeson, L. Carpenter, L. Casten, L. Coppola, L. Cordiero, L. DeMarco, L. D. Pacheco, L. F. Corzo, L. H. Shulman, L. K. Walsh, L. Lesher, L. M. Herbert, L. M. Prock, L. Malloch, L. Mann, L. P. Grosvenor, L. Simon, L. V. Soorya, L. Wasserburg, L. Yeh, L. Y. Huang-Storms, M. Alessandri, M. A. Popp, M. Baer, M. Beckwith, M. Casseus, M. Coughlin, M. Currin, M. Cutri, M. D. Mallardi, M. DuBois, M. Dunlevy, M. E. Butler, M. Frayne, M. F. Gwynette, M. Ghaziuddin, M. Haley, M. Heyman, M. Hojlo, M. Jordy, M. J. Morrier, M. Kowanda, M. Koza, M. Lopez, M. McTaggart, M. Norris, M. N. Hale, M. O’Neil, M. Printen, M. Rayos, M. Sabiha, M. Sahin, M. Sarris, M. Shir, M. Siegel, M. Steele, M. Sweeney, M. Tafolla, M. Valicenti-McDermott, M. Verdi, M. Y. Dennis, N. Alvarez, N. Bardett, N. Berger, N. Calderon, N. Decius, N. Gonzalez, N. Harris, N. Lawson, N. Lillie, N. Lo, N. Long, N. M. Russo-Ponsaran, N. Madi, N. Mccoy, N. Nagpal, N. Rodriguez, N. Russell, N. Shah, N. Takahashi, N. Targalia, O. Newman, O. Y. Ousley, P. Heydemann, P. Manning, P. S. Carbone, R. A. Bernier, R. A. Gordon, R. C. Shaffer, R. D. Annett, R. D. Clark, R. Jou, R. J. Landa, R. K. Earl, R. Libove, R. Marini, R. N. Doan, R. P. Goin-Kochel, R. Rana, R. Remington, R. Shikov, R. T. Schultz, S. Aberle, S. Birdwell, S. Boland, S. Booker, S. Carpenter, S. Chintalapalli, S. Conyers, S. D’Ambrosi, S. Eldred, S. Francis, S. Ganesan, S. Hepburn, S. Horner, S. Hunter, S. J. Brewster, S. J. Lee, S. Jacob, S. Jean, S. Hyun, S. Kramer, S. L. Friedman, S. Licona, S. Littlefield, S. M. Kanne, S. Mastel, S. Mathai, S. Melnyk, S. Michaels, S. Mohiuddin, S. Palmer, S. Plate, S. Qiu, S. Randall, S. Sandhu, S. Santangelo, S. Shah, S. Skinner, S. Thompson, S. White, S. White, S. Xiao, S. Xu, S. Xu, T. Chen, T. Greene, T. Ho, T. Ibanez, T. Koomar, T. Pramparo, T. Rutter, T. Shaikh, T. Tran, T. W. Yu, V. Galbraith, V. Gazestani, V. J. Myers, V. Ranganathan, V. Singh, W. C. Weaver, W. CaI, W. Chin, W. S. Yang, Y. B. Choi, Z. E. Warren, J. J. Michaelson, N. Volfovsky, E. E. Eichler, Y. Shen, W. K. Chung, Integrating de novo and inherited variants in 42,607 autism cases identifies mutations in new moderate-risk genes. Nat Genet 54, 1305–1319 (2022).

74. P. Xia, L. Zhou, J. Guan, W. Ding, Y. Liu, Splicing factor PRP-19 regulates mitochondrial stress response. Life Metabolism 1, 81–93 (2022).

75. C. J. Jeffery, Protein moonlighting: what is it, and why is it important? Phil. Trans. R. Soc. B 373, 20160523 (2018).

76. D. H. E. W. Huberts, I. J. Van Der Klei, Moonlighting proteins: An intriguing mode of multitasking. Biochimica et Biophysica Acta (BBA) - Molecular Cell Research 1803, 520– 525 (2010).

77. B. Henderson, A. C. R. Martin, Protein moonlighting: a new factor in biology and medicine. Biochemical Society Transactions 42, 1671–1678 (2014).

78. A. M. Nargund, M. W. Pellegrino, C. J. Fiorese, B. M. Baker, C. M. Haynes, Mitochondrial Import Efficiency of ATFS-1 Regulates Mitochondrial UPR Activation. Science 337, 587– 590 (2012).

79. C. M. Haynes, Y. Yang, S. P. Blais, T. A. Neubert, D. Ron, The Matrix Peptide Exporter HAF-1 Signals a Mitochondrial UPR by Activating the Transcription Factor ZC376.7 in C. elegans. Molecular Cell 37, 529–540 (2010).

80. C. Münch, J. W. Harper, Mitochondrial unfolded protein response controls matrix pre-RNA processing and translation. Nature 534, 710–713 (2016).

81. C. J. Fiorese, A. M. Schulz, Y.-F. Lin, N. Rosin, M. W. Pellegrino, C. M. Haynes, The Transcription Factor ATF5 Mediates a Mammalian Mitochondrial UPR. Current Biology 26, 2037–2043 (2016).

82. P. M. Quirós, M. A. Prado, N. Zamboni, D. D’Amico, R. W. Williams, D. Finley, S. P. Gygi, J. Auwerx, Multi-omics analysis identifies ATF4 as a key regulator of the mitochondrial stress response in mammals. Journal of Cell Biology 216, 2027–2045 (2017).

83. R. H. Houtkooper, L. Mouchiroud, D. Ryu, N. Moullan, E. Katsyuba, G. Knott, R. W. Williams, J. Auwerx, Mitonuclear protein imbalance as a conserved longevity mechanism. Nature 497, 451–457 (2013).

84. K. Pakos-Zebrucka, I. Koryga, K. Mnich, M. Ljujic, A. Samali, A. M. Gorman, The integrated stress response. EMBO Reports 17, 1374–1395 (2016).

85. M. Costa-Mattioli, P. Walter, The integrated stress response: From mechanism to disease. Science 368, eaat5314 (2020).

86. X. Guo, G. Aviles, Y. Liu, R. Tian, B. A. Unger, Y.-H. T. Lin, A. P. Wiita, K. Xu, M. A. Correia, M. Kampmann, Mitochondrial stress is relayed to the cytosol by an OMA1– DELE1–HRI pathway. Nature 579, 427–432 (2020).

87. E. Fessler, E.-M. Eckl, S. Schmitt, I. A. Mancilla, M. F. Meyer-Bender, M. Hanf, J. Philippou-Massier, S. Krebs, H. Zischka, L. T. Jae, A pathway coordinated by DELE1 relays mitochondrial stress to the cytosol. Nature 579, 433–437 (2020).

88. E. Mick, D. V. Titov, O. S. Skinner, R. Sharma, A. A. Jourdain, V. K. Mootha, Distinct mitochondrial defects trigger the integrated stress response depending on the metabolic state of the cell. eLife 9, e49178 (2020).

89. Min, S., et al., Mitoribosomal Deregulation Drives Senescence via TPP1-Mediated Telomere Deprotection. Cells, 2022. 11(13).

90. J. G. Doench, N. Fusi, M. Sullender, M. Hegde, E. W. Vaimberg, K. F. Donovan, I. Smith, Z. Tothova, C. Wilen, R. Orchard, H. W. Virgin, J. Listgarten, D. E. Root, Optimized sgRNA design to maximize activity and minimize off-target effects of CRISPR-Cas9. Nat Biotechnol 34, 184–191 (2016).

91. K. R. Sanson, R. E. Hanna, M. Hegde, K. F. Donovan, C. Strand, M. E. Sullender, E. W. Vaimberg, A. Goodale, D. E. Root, F. Piccioni, J. G. Doench, Optimized libraries for CRISPR-Cas9 genetic screens with multiple modalities. Nat Commun 9, 5416 (2018).

92. F. A. Ran, P. D. Hsu, J. Wright, V. Agarwala, D. A. Scott, F. Zhang, Genome engineering using the CRISPR-Cas9 system. Nat Protoc 8, 2281–2308 (2013).

93. Khawaja, A., et al., Insights into mitoribosomal biogenesis from recent structural studies. Trends Biochem Sci, 2023. 48(7): p. 629–641.

94. Lavdovskaia, E., et al., A roadmap for ribosome assembly in human mitochondria. Nat Struct Mol Biol, 2024. 31(12): p. 1898–1908.

95. B. Ruzzenente, M. D. Metodiev, “Linear Density Sucrose Gradients to Study Mitoribosomal Biogenesis in Tissue-Specific Knockout Mice” in Mouse Genetics, S. R. Singh, R. M. Hoffman, A. Singh, Eds. (Springer US, New York, NY, 2021; http://link.springer.com/10.1007/978-1-0716-1008-4_3)vol. 2224 of Methods in Molecular Biology, pp. 47–60.

96. R. J. A. Wanders, H. R. Waterham, S. Ferdinandusse, Metabolic Interplay between Peroxisomes and Other Subcellular Organelles Including Mitochondria and the Endoplasmic Reticulum. Front. Cell Dev. Biol. 3 (2016).

97. A. Koch, Y. Yoon, N. A. Bonekamp, M. A. McNiven, M. Schrader, A Role for Fis1 in Both Mitochondrial and Peroxisomal Fission in Mammalian Cells□D.

98. S. Gandre-Babbe, A. M. Van Der Bliek, The Novel Tail-anchored Membrane Protein Mff Controls Mitochondrial and Peroxisomal Fission in Mammalian Cells. MBoC 19, 2402–2412 (2008).

99. I. G. Castro, D. M. Richards, J. Metz, J. L. Costello, J. B. Passmore, T. A. Schrader, A. Gouveia, D. Ribeiro, M. Schrader, A role for Mitochondrial Rho GTPase 1 (MIRO1) in motility and membrane dynamics of peroxisomes. Traffic 19, 229–242 (2018).

100. N. E. Braverman, G. V. Raymond, W. B. Rizzo, A. B. Moser, M. E. Wilkinson, E. M. Stone, S. J. Steinberg, M. F. Wangler, E. T. Rush, J. G. Hacia, M. Bose, Peroxisome biogenesis disorders in the Zellweger spectrum: An overview of current diagnosis, clinical manifestations, and treatment guidelines. Molecular Genetics and Metabolism 117, 313–321 (2016).

101. J. Wegrzyn, R. Potla, Y.-J. Chwae, N. B. V. Sepuri, Q. Zhang, T. Koeck, M. Derecka, K. Szczepanek, M. Szelag, A. Gornicka, A. Moh, S. Moghaddas, Q. Chen, S. Bobbili, J. Cichy, J. Dulak, D. P. Baker, A. Wolfman, D. Stuehr, M. O. Hassan, X.-Y. Fu, N. Avadhani, J. I. Drake, P. Fawcett, E. J. Lesnefsky, A. C. Larner, Function of Mitochondrial Stat3 in Cellular Respiration. Science 323, 793–797 (2009).

102. N. D. Marchenko, A. Zaika, U. M. Moll, Death Signal-induced Localization of p53 Protein to Mitochondria. Journal of Biological Chemistry 275, 16202–16212 (2000).

103. V. D. Antonenkov, S. Grunau, S. Ohlmeier, J. K. Hiltunen, Peroxisomes Are Oxidative Organelles. Antioxidants & Redox Signaling 13, 525–537 (2010).

104. M. Fransen, M. Nordgren, B. Wang, O. Apanasets, Role of peroxisomes in ROS/RNS-metabolism: Implications for human disease. Biochimica et Biophysica Acta (BBA) - Molecular Basis of Disease 1822, 1363–1373 (2012).

105. C. Lismont, M. Nordgren, P. P. Van Veldhoven, M. Fransen, Redox interplay between mitochondria and peroxisomes. Front. Cell Dev. Biol. 3 (2015).

106. C. Lismont, I. Revenco, M. Fransen, Peroxisomal Hydrogen Peroxide Metabolism and Signaling in Health and Disease. IJMS 20, 3673 (2019).

107. S. Kemp, R. J. A. Wanders, X-linked adrenoleukodystrophy: Very long-chain fatty acid metabolism, ABC half-transporters and the complicated route to treatment. Molecular Genetics and Metabolism 90, 268–276 (2007).

108. K. Itoh, N. Wakabayashi, Y. Katoh, T. Ishii, K. Igarashi, J. D. Engel, M. Yamamoto, Keap1 represses nuclear activation of antioxidant responsive elements by Nrf2 through binding to the amino-terminal Neh2 domain. Genes Dev. 13, 76–86 (1999).

109. A. Reyes, P. Favia, S. Vidoni, V. Petruzzella, M. Zeviani, RCC1L (WBSCR16) isoforms coordinate mitochondrial ribosome assembly through their interaction with GTPases. PLoS Genet 16, e1008923 (2020).

110. S. Zhang, Z. Dong, Y. Feng, W. Guo, C. Zhang, Y. Shi, Z. Zhao, J. Wang, G. Ning, G. Huang, WBSCR16 is essential for mitochondrial 16S rRNA processing in mammals. Nucleic Acids Research 53, gkae1325 (2025).

111. Y. Cámara, J. Asin-Cayuela, C. B. Park, M. D. Metodiev, Y. Shi, B. Ruzzenente, C. Kukat, B. Habermann, R. Wibom, K. Hultenby, T. Franz, H. Erdjument-Bromage, P. Tempst, B. M. Hallberg, C. M. Gustafsson, N.-G. Larsson, MTERF4 Regulates Translation by Targeting the Methyltransferase NSUN4 to the Mammalian Mitochondrial Ribosome. Cell Metabolism 13, 527–539 (2011).

112. M. Lagouge, A. Mourier, H. J. Lee, H. Spåhr, T. Wai, C. Kukat, E. Silva Ramos, E. Motori, J. D. Busch, S. Siira, German Mouse Clinic Consortium, E. Kremmer, A. Filipovska, N.-G. Larsson, SLIRP Regulates the Rate of Mitochondrial Protein Synthesis and Protects LRPPRC from Degradation. PLoS Genet 11, e1005423 (2015).

113. H. Antonicka, F. Sasarman, T. Nishimura, V. Paupe, E. A. Shoubridge, The Mitochondrial RNA-Binding Protein GRSF1 Localizes to RNA Granules and Is Required for Posttranscriptional Mitochondrial Gene Expression. Cell Metabolism 17, 386–398 (2013).

114. A. A. Jourdain, M. Koppen, C. D. Rodley, K. Maundrell, N. Gueguen, P. Reynier, A. M. Guaras, J. A. Enriquez, P. Anderson, M. Simarro, J.-C. Martinou, A Mitochondria-Specific Isoform of FASTK Is Present In Mitochondrial RNA Granules and Regulates Gene Expression and Function. Cell Reports 10, 1110–1121 (2015).

115. H. P. Harding, Y. Zhang, A. Bertolotti, H. Zeng, D. Ron, Perk Is Essential for Translational Regulation and Cell Survival during the Unfolded Protein Response. Molecular Cell 5, 897– 904 (2000).

116. Walter P, Ron D. The unfolded protein response: from stress pathway to homeostatic regulation. Science. 2011 Nov 25;334(6059):1081–6. doi: 10.1126/science.1209038. PMID: 22116877.

117. D. A. Bota, K. J. A. Davies, Lon protease preferentially degrades oxidized mitochondrial aconitase by an ATP-stimulated mechanism. Nat Cell Biol 4, 674–680 (2002).

118. P. M. Quirós, Y. Español, R. Acín-Pérez, F. Rodríguez, C. Bárcena, K. Watanabe, E. Calvo, M. Loureiro, M. S. Fernández-García, A. Fueyo, J. Vázquez, J. A. Enríquez, C. López-Otín, ATP-Dependent Lon Protease Controls Tumor Bioenergetics by Reprogramming Mitochondrial Activity. Cell Reports 8, 542–556 (2014).

119. J. R. Warner, The economics of ribosome biosynthesis in yeast. Trends in Biochemical Sciences 24, 437–440 (1999).

120. E. Thomson, S. Ferreira-Cerca, E. Hurt, Eukaryotic ribosome biogenesis at a glance. Journal of Cell Science 126, 4815–4821 (2013).

121. E. Metzl-Raz, M. Kafri, G. Yaakov, I. Soifer, Y. Gurvich, N. Barkai, Principles of cellular resource allocation revealed by condition-dependent proteome profiling. eLife 6, e28034 (2017).

122. A. Peeters, A. B. Shinde, R. Dirkx, J. Smet, K. De Bock, M. Espeel, I. Vanhorebeek, A. Vanlander, R. Van Coster, P. Carmeliet, M. Fransen, P. P. Van Veldhoven, M. Baes, Mitochondria in peroxisome-deficient hepatocytes exhibit impaired respiration, depleted DNA, and PGC-1α independent proliferation. Biochimica et Biophysica Acta (BBA) - Molecular Cell Research 1853, 285–298 (2015).

123. T. Powers, P. Walter, Regulation of Ribosome Biogenesis by the Rapamycin-sensitive TOR-signaling Pathway in *Saccharomyces cerevisiae*. MBoC 10, 987–1000 (1999).

124. C. Mayer, I. Grummt, Ribosome biogenesis and cell growth: mTOR coordinates transcription by all three classes of nuclear RNA polymerases. Oncogene 25, 6384–6391 (2006).

125. R. A. Saxton, D. M. Sabatini, mTOR Signaling in Growth, Metabolism, and Disease. Cell 168, 960–976 (2017).

126. R. Dirkx, I. Vanhorebeek, K. Martens, A. Schad, M. Grabenbauer, D. Fahimi, P. Declercq, P. P. Van Veldhoven, M. Baes, Absence of peroxisomes in mouse hepatocytes causes mitochondrial and ER abnormalities†. Hepatology 41, 868–878 (2005).

127. E. Nuebel, J. T. Morgan, S. Fogarty, J. M. Winter, S. Lettlova, J. A. Berg, Y.-C. Chen, C. U. Kidwell, J. A. Maschek, K. J. Clowers, C. Argyriou, L. Chen, I. Wittig, J. E. Cox, M. Roh-Johnson, N. Braverman, J. Bonkowsky, S. P. Gygi, J. Rutter, The biochemical basis of mitochondrial dysfunction in Zellweger Spectrum Disorder. EMBO Rep 22, EMBR202051991 (2021).

128. J. E. Legakis, J. I. Koepke, C. Jedeszko, F. Barlaskar, L. J. Terlecky, H. J. Edwards, P. A. Walton, S. R. Terlecky, Peroxisome Senescence in Human Fibroblasts. MBoC 13, 4243–4255 (2002).

129. J. I. Koepke, K. Nakrieko, C. S. Wood, K. K. Boucher, L. J. Terlecky, P. A. Walton, S. R. Terlecky, Restoration of Peroxisomal Catalase Import in a Model of Human Cellular Aging. Traffic 8, 1590–1600 (2007).

130. V. I. Titorenko, S. R. Terlecky, Peroxisome Metabolism and Cellular Aging. Traffic 12, 252– 259 (2011).

131. A. Vercaemst, M. Zhao, R. Chai, C. Lismont, M. Fransen, Peroxisomes in Aging: Guardians of Cellular Resilience and Function. Cells 15, 254 (2026).

132. J. Campisi, F. d’Adda Di Fagagna, Cellular senescence: when bad things happen to good cells. Nat Rev Mol Cell Biol 8, 729–740 (2007).

133. J.-P. Coppé, C. K. Patil, F. Rodier, Y. Sun, D. P. Muñoz, J. Goldstein, P. S. Nelson, P.-Y. Desprez, J. Campisi, Senescence-Associated Secretory Phenotypes Reveal Cell-Nonautonomous Functions of Oncogenic RAS and the p53 Tumor Suppressor. PLoS Biol 6, e301 (2008).

134. J. C. Acosta, A. O’Loghlen, A. Banito, M. V. Guijarro, A. Augert, S. Raguz, M. Fumagalli, M. Da Costa, C. Brown, N. Popov, Y. Takatsu, J. Melamed, F. d’Adda Di Fagagna, D. Bernard, E. Hernando, J. Gil, Chemokine Signaling via the CXCR2 Receptor Reinforces Senescence. Cell 133, 1006–1018 (2008).

135. C. D. Wiley, J. Campisi, The metabolic roots of senescence: mechanisms and opportunities for intervention. Nat Metab 3, 1290–1301 (2021).

136. W. Huang, L. J. Hickson, A. Eirin, J. L. Kirkland, L. O. Lerman, Cellular senescence: the good, the bad and the unknown. Nat Rev Nephrol 18, 611–627 (2022).

137. S. Victorelli, H. Salmonowicz, J. Chapman, H. Martini, M. G. Vizioli, J. S. Riley, C. Cloix, E. Hall-Younger, J. Machado Espindola-Netto, D. Jurk, A. B. Lagnado, L. Sales Gomez, J. N. Farr, D. Saul, R. Reed, G. Kelly, M. Eppard, L. C. Greaves, Z. Dou, N. Pirius, K. Szczepanowska, R. A. Porritt, H. Huang, T. Y. Huang, D. A. Mann, C. A. Masuda, S. Khosla, H. Dai, S. H. Kaufmann, E. Zacharioudakis, E. Gavathiotis, N. K. LeBrasseur, X. Lei, A. G. Sainz, V. I. Korolchuk, P. D. Adams, G. S. Shadel, S. W. G. Tait, J. F. Passos, Apoptotic stress causes mtDNA release during senescence and drives the SASP. Nature 622, 627–636 (2023).

138. B. Wang, J. Han, J. H. Elisseeff, M. Demaria, The senescence-associated secretory phenotype and its physiological and pathological implications. Nat Rev Mol Cell Biol 25, 958–978 (2024).

139. T. Natsume, T. Kiyomitsu, Y. Saga, M. T. Kanemaki, Rapid Protein Depletion in Human Cells by Auxin-Inducible Degron Tagging with Short Homology Donors. Cell Reports 15, 210–218 (2016).

140. W. Li, H. Xu, T. Xiao, L. Cong, M. I. Love, F. Zhang, R. A. Irizarry, J. S. Liu, M. Brown, X. S. Liu, MAGeCK enables robust identification of essential genes from genome-scale CRISPR/Cas9 knockout screens.

141. M. Spinazzi, A. Casarin, V. Pertegato, L. Salviati, C. Angelini, Assessment of mitochondrial respiratory chain enzymatic activities on tissues and cultured cells. Nat Protoc 7, 1235–1246 (2012).

142. M. D. Metodiev, N. Lesko, C. B. Park, Y. Cámara, Y. Shi, R. Wibom, K. Hultenby, C. M. Gustafsson, N.-G. Larsson, Methylation of 12S rRNA Is Necessary for In Vivo Stability of the Small Subunit of the Mammalian Mitochondrial Ribosome. Cell Metabolism 9, 386–397 (2009).

